# Genomic and biochemical contexts determine the physiological role of a horizontally acquired gene

**DOI:** 10.64898/2026.07.09.737255

**Authors:** Roberto E. Bruna, Alagu Lakshmi Selvaraj, Somok Bhowmik, Christopher G. Kendra, Ryan Heister, Mauricio H. Pontes

## Abstract

The horizontally acquired *mgtC* gene from *Salmonella enterica* confers this bacterium the abilities to survive episodes of magnesium (Mg^2+^) starvation, and to replicate in mammalian macrophages. The former property allows bacteria to persist in the environment through periods of Mg^2+^ depletion, whereas the latter allows *S. enterica* to overcome self-limiting intestinal colonization and cause an invasive systemic infection in susceptible mammalian hosts. Even though the biochemical function of MgtC is not completely understood, this protein is thought to function primarily by preventing the production of toxic levels of Mg^2+^-chelating adenosine triphosphate (ATP). In the current work, we investigated the physiological roles of *mgtC* homologs from an array of bacterial species, by probing the processes controlled by this gene during replication in low Mg^2+^ medium and in macrophages. We determined that MgtC homologs that do not participate in Pi homeostasis during Mg^2+^ starvation and do not promote intramacrophage replication in their resident species can partake in these processes when expressed in *S. enterica*. This indicates that the function of this protein is context dependent. Accordingly, we show that the physiological processes affected by *S. enterica* MgtC vary, depending on whether the bacteria replicate in low Mg^2+^ medium or inside macrophages. While these results suggest that MgtC is a regulator, they also demonstrate that horizontally acquired genes can assume different roles, depending on the genome and the biochemical context into which they are inserted.

**Importance:** The *mgtC* gene encodes an inner membrane protein that has been horizontally acquired by multiple bacterial species, including several mammalian pathogens. In *Salmonella enterica*, MgtC promotes replication in mammalian macrophages and allows this bacterium to survive cytoplasmic magnesium (Mg^2+^) starvation. These phenotypes are thought to result from MgtC’s inhibition of Pi metabolism and ATP production, which prevents the accumulation of toxic levels of Mg^2+^-chelating ATP and disrupts other physiological processes that are strictly dependent on Mg^2+^, such as ribosome assembly and translation. In the current study, we show that processes that are controlled by MgtC vary with the genetic and biochemical contexts in which this protein is expressed. While establishing a broader role for MgtC as a regulator, our findings illustrate how horizontally acquired regulatory genes can potentiate regulatory interactions, facilitating the evolution of new traits.

## Introduction

The acquisition of genetic information from exogenous sources is a major driving force in bacterial evolution. Incoming genes that are incorporated and maintained in bacterial lineages often confer adaptive traits to recipient cells; enhancing their fitness in the current niche, enabling ecological niche expansion and promoting adaptive radiation [1–5]. In this process, new traits can surface immediately, solely through the expression of information contained within horizontally acquired genes. For example, a lineage can become resistant to an antibiotic after obtaining and expressing genes encoding resistant determinants to that antibiotic [6]. New properties can also emerge gradually, as foreign genes are integrated into new regulatory networks and biochemical surroundings. For instance, the horizontal acquisition and progressive integration of a gene encoding a regulator can enable the evolution of regulatory interactions that alter the ancestral biochemical environment within a lineage, thereby giving rise to new traits upon which selection can act [7–19].

The *mgtC* gene has been horizontally acquired by multiple bacterial species, including several major human pathogens [20] (Fig. 1). This gene encodes an inner membrane protein that is typically required for bacterial survival to cytoplasmic magnesium (Mg^2+^) starvation and virulence [21–44]. In *Salmonella enterica*, *mgtC* is located within the horizontally acquired chromosomal region designated *Salmonella* Pathogenicity Island 3. It is expressed when bacteria replicate within mammalian macrophages or experience cytoplasmic Mg^2+^ starvation during growth in defined laboratory medium [21, 22, 45–50]. Inactivating mutations in *mgtC* abolish intramacrophage replication, thereby severely attenuating *S. enterica* virulence in a mouse model of systemic infection [22, 24, 31, 51]. Likewise, during cytoplasmic Mg^2+^ starvation, these mutations halt growth, causing a gradual loss of viability and a myriad of accompanying phenotypes. These include impaired cellular septation, overproduction of a biofilm extracellular matrix component, inhibition of ribosomal subunit assembly and thus translation, and a substantial increase in the importation and assimilation of phosphate (PO_2_^-3^, Pi) into adenosine triphosphate (ATP), which alters downstream processes such as catabolite repression [22, 25, 30, 45, 52–56].

**Figure 1.**
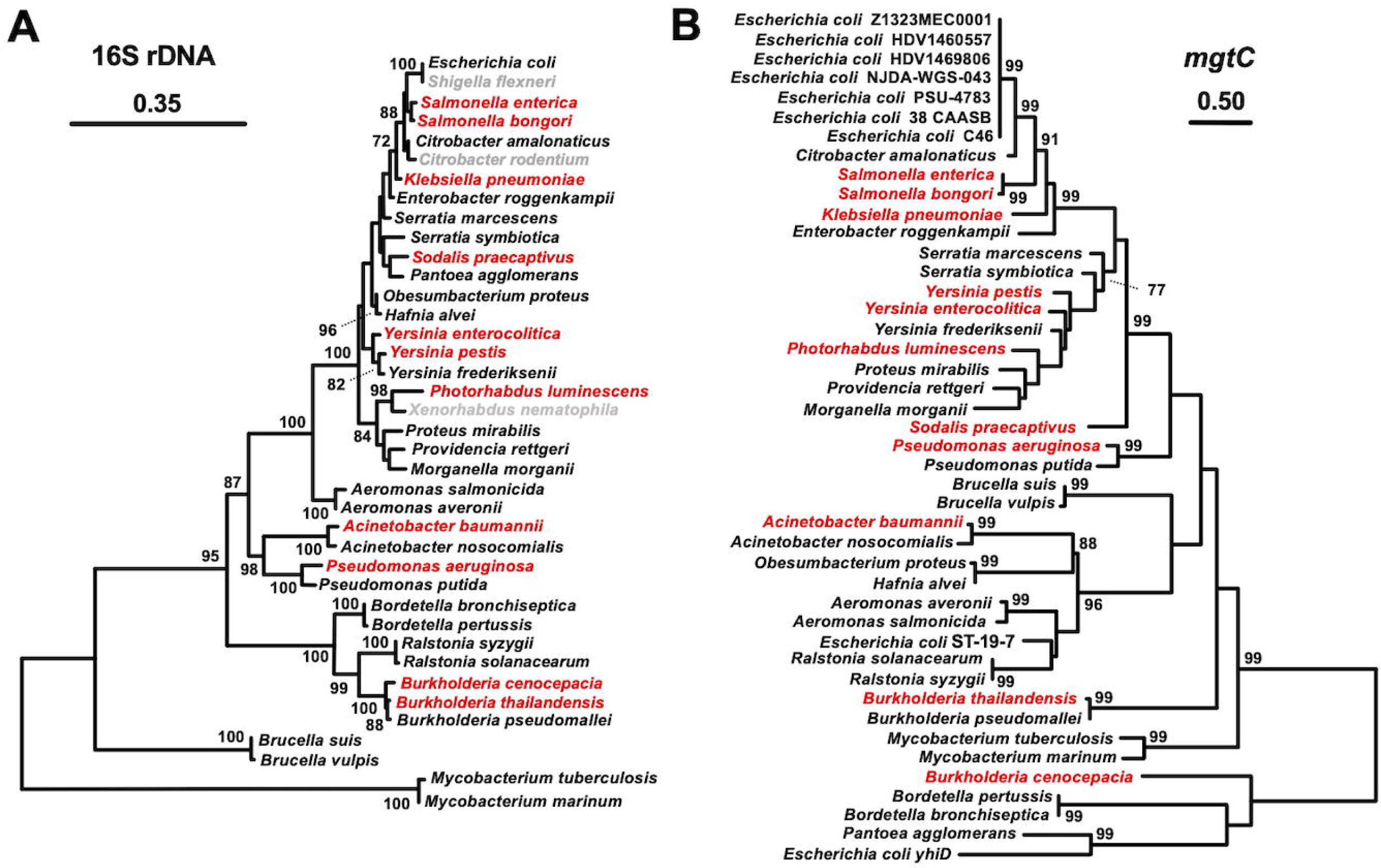
Phylogenies of sequences from selected bacterial species based on maximum likelihood analyses of 16S rRNA (left-hand side tree) and *mgtC*/*yhiD* homologs (right-hand side tree). **(A)** Maximum likelihood 16S rRNA phylogeny of selected species. Note that sequenced representatives of *Shigella flexneri*, *Citrobacter rodentium* and *Xenorhabdus nematophila* (in gray) lack *mgtC* homologs. **(B)** Maximum likelihood codon-aligned *mgtC*/*yhiD* phylogeny of selected species. Note that the presence of *mgtC* is sporadic within certain genera (e.g., *Citrobacter rodentium* lacks *mgtC*, while *Citrobacter amalonaticus* encodes a *mgtC* homolog) and species (e.g., while most sequenced *E. coli* strains lack a *mgtC*, some, such as Z1323MEC0001, PSU-4683, and ST-19-7, have horizontally acquired this gene). Topological incongruency between (A) 16S rRNA and (B) *mgtC*/*yhiD* phylogenies at deeper nodes, with closely related species harboring distantly related *mgtC* homologs, indicates horizontal gene acquisition. For (A) and (B): Bacterial species and their respective *mgtC* homologs investigated in this study are depicted in red. Maximum-likelihood bootstrap values with bootstrap support >70% are shown adjacent to nodes in the trees.

Several of these phenotypes, including growth arrest and loss of viability, can be suppressed by decreasing intracellular ATP, thus supporting the notion that the physiological role of MgtC is to maintain ATP homeostasis [25, 52–54, 57]. Consequently, the phenotypic effects of *mgtC* inactivation are posited to result from the depletion of free cytoplasmic Mg^2+^ by the formation of ATP:Mg^2+^ salt, and the disruption of cellular processes that are strictly dependent on Mg^2+^. That is, MgtC increases the concentration of free intracellular Mg^2+^ by preventing the overproduction of ATP [58, 59]. However, two main lines of evidence suggest that the physiological function of this protein depends on the biochemical context in which it is expressed. First, a mutation that alters a single amino acid residue in MgtC hinders intramacrophage replication without affecting survival under Mg^2+^ starvation. In other words, the phenotypes conferred by MgtC within macrophages and during Mg^2+^ starvation can be genetically separated [30]. Second, *S. enterica* requires MgtC for intramacrophage replication, regardless of the macrophage’s ability to restrict the access of internalized bacteria to Mg^2+^. In other words, *S. enterica* requires MgtC even in macrophages that are abundant in Mg^2+^ [31, 60].

In the current study, we examined the impact of context on MgtC function. We probed the physiological roles of *mgtC* homologs from species spanning a wide phylogenetic distribution (Fig. 1). We determined that most homologs promote growth during Mg^2+^ starvation by reducing Pi intake and ATP synthesis. Surprisingly, MgtC homologs that no longer promote these traits in their current species can inhibit ATP synthesis and promote growth of a *S. enterica mgtC* mutant during cytoplasmic Mg^2+^ starvation. Likewise, we establish that MgtC homologs that do not affect intramacrophage replication in their present species, can promote replication of a *S. enterica mgtC* mutant within these eukaryotic cells. Finally, we demonstrate that the roles of *S. enterica* MgtC on Pi metabolism and translation homeostasis that take place during cytoplasmic Mg^2+^ starvation [52–54] do not occur during intramacrophage replication. This suggests that these processes result from interactions between MgtC and other cellular components, rather than from intrinsic properties of this protein. These results also indicate that the role of MgtC on bacterial physiology depends on the genomic content and ultimately, on the biochemical environment in which this protein is integrated following horizontal acquisition.

## Results

### MgtC control of phosphate and ATP homeostasis during cytoplasmic Mg^2+^ stress is conserved and widespread

When *S. enterica* is inoculated in defined medium containing low (0.01 mM) Mg^2+^, cells grow rapidly for the first 3-4 h by consuming most of the Mg^2+^ available in the medium using the housekeeping Mg^2+^ transporter CorA. Subsequently, CorA ceases to efficiently transport Mg^2+^ and the cells experience a decrease in free cytoplasmic Mg^2+^. This decrease in free cytoplasmic Mg^2+^ slows translation, thereby decreasing ATP consumption. While the accumulation of ATP and its chelation of free Mg^2+^ exacerbate the stress, the decreased translation promotes MgtC expression [49, 50, 52, 58]. MgtC, in turn, inhibits Pi acquisition, hindering ATP synthesis, thereby freeing cytoplasmic Mg^2+^ and restoring bacterial growth [54, 57].

While, to date, all *mgtC* homologs have been shown to promote bacterial growth during Mg^2+^ starvation [22, 26, 28–30, 33–36, 40–43, 45], we sought to test whether they also inhibit Pi transport and ATP synthesis to promote bacterial growth. To this end, we examined the properties conferred by MgtC homologs in ten bacterial species: *Salmonella bongori*, *Klebsiella pneumoniae*, *Yersinia pestis*, *Yersinia enterocolitica*, *Photorhabdus luminescens*, *Sodalis praecaptivus*, *Acinetobacter baumannii*, *Burkholderia thailandensis*, *Burkholderia cenocepacia* and *Pseudomonas aeruginosa*. These species inhabit diverse niches, span a wide phylogenetic distribution (Fig. 1A) and encode *mgtC* homologs from distinct lineages that vary in sequence divergence and the timing of horizontal acquisition (Fig. 1B).

We determined that *mgtC* inactivation hindered the growth of *S. bongori*, *K. pneumoniae*, *Y. pestis*, *Y. enterocolitica*, *A. baumannii*, *B. cenocepacia* [33], *B. thailandensis* and *P. aeruginosa* [36] (Fig. 2A). The growth defect of these *mgtC* mutant strains was improved by removing exogenous Pi from the medium. However, the toxic effect of exogenous Pi removal on growth varied across species. Whereas Pi limitation only partially rescued the growth of the *Y. pestis mgtC* mutant, it completely restored the growth of *S. bongori mgtC* to wild-type levels (Fig. 2A). Interestingly, for *K. pneumoniae*, *P. aeruginosa*, *B. cenocepacia* and, *B. thailandensis*, Pi limitation improved the growth of both wild-type and *mgtC* mutant strains (Fig. 2A). This later phenomenon was also observed in *Escherichia coli*, likely reflecting the fact that Pi is acutely toxic to cells experiencing cytoplasmic Mg^2+^ starvation [53, 54]. However, consistent with previous results [30, 52, 53], inactivation of *E. coli*’s *yhiD* gene, which encodes the protein with the highest level of amino acid sequence identity to the MgtC from *S. enterica*, did not affect growth during Mg^2+^ limitation (Fig. 2A).

**Figure 2.**
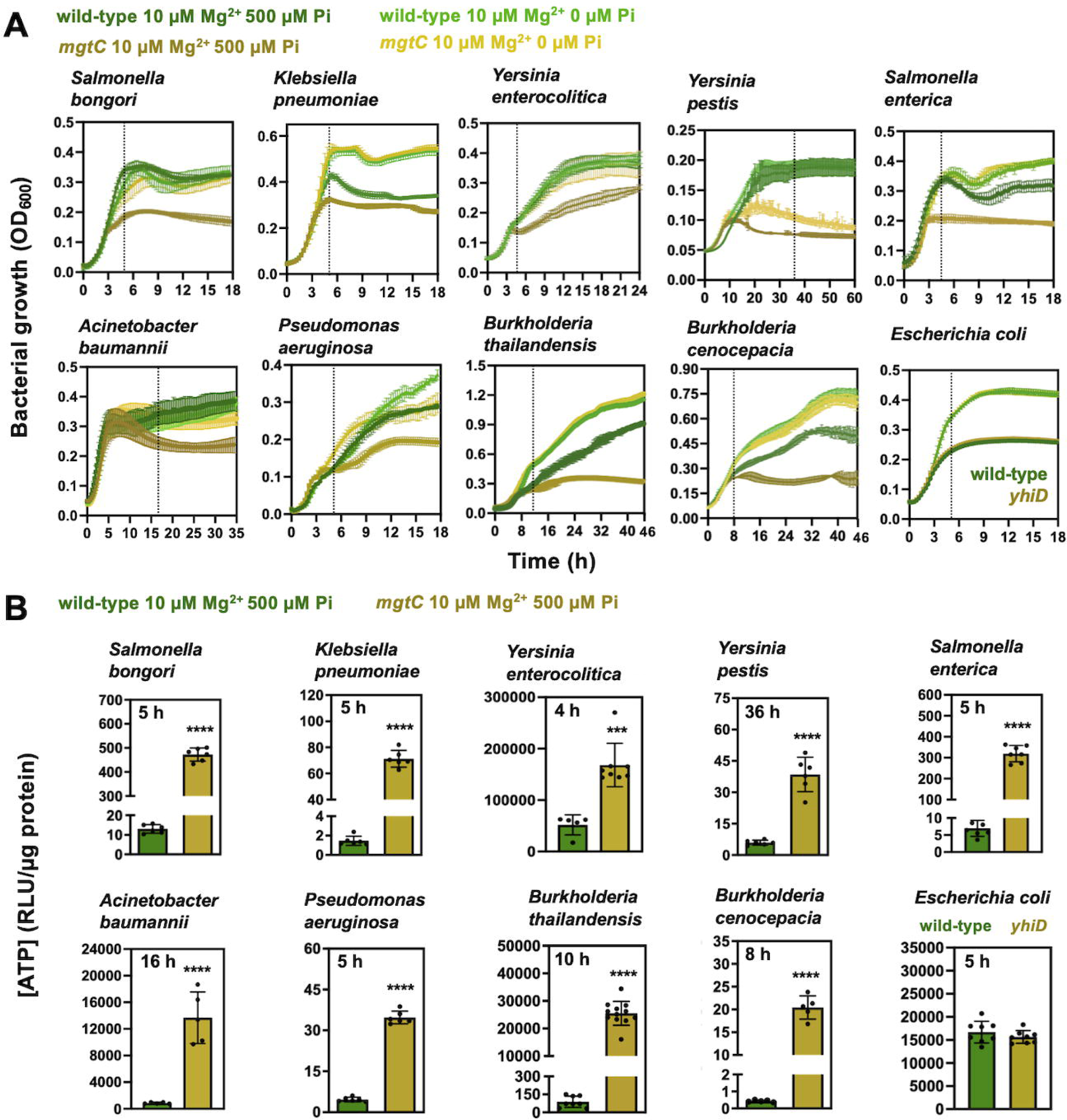
Role of MgtC homologs in controlling phosphate metabolism and ATP production during magnesium starvation. **(A)** Growth curves of wild-type and isogenic *mgtC* mutant strains of selected bacterial species in MOPS medium containing 10 µM MgCl_2_ and either 500 µM or no exogenous phosphate in the form of K_2_HPO_4_ (note that casamino acid mixture present in the media contains approximately 165 μM of Pi and supports bacterial growth). Dashed lines indicate time points when intracellular ATP levels, displayed in (B), were measured. **(B)** Relative intracellular ATP levels of strains depicted in (A). ATP measurements were conducted in MOPS medium containing 10 µM MgCl_2_ and 500 µM of K_2_HPO_4_ at the indicated time points. ***P < 0.001 and ****P < 0.0001, unpaired t-tests between ATP concentrations in populations of wild-type and *mgtC* mutant strains. For all graphs means ± SDs of at least three biological replicates are shown. Graphs depict representative data from one of at least three independent experiments. Measurements were conducted in the following strains: *A. baumannii* wild-type (ATCC 17978) and *mgtC* (A1S_2069), *B. cenocepacia* wild-type (K56-2) and *mgtC* (KEM1); B*. thailandensis* wild-type (E264) and *mgtC* (MP1936); *E. coli* wild-type (MG1655) and *yhiD* (MP751); *K. pneumoniae* wild-type (MKP103) and *mgtC* (KP5811); *P. aeruginosa* wild-type (PAO1) and *mgtC* (PW8814); *S. bongori* wild-type (S3041) and *mgtC* (RB52); *S. enterica* wild-type (14028s) and *mgtC* (EL4); *Y. enterocolitica* wild-type (JB580v) *mgtC* (MP2500); *Y. pestis* wild-type (KIM6) and *mgtC* (RB51).

Subsequently, we measured the ATP concentrations in wild-type and isogenic *mgtC* mutants after the onset of growth defects induced by cytoplasmic Mg^2+^-starvation. We established that *mgtC* inactivation led to significant increases in ATP concentrations in *S. bongori*, *K. pneumoniae*, *Y. pestis*, *Y. enterocolitica*, *A. baumannii*, *B. cenocepacia*, *B. thailandensis* and *P. aeruginosa* indicating that the MgtC proteins of these species behave like that of *S. enterica* [53, 54] (Fig. 2B). As expected, inactivation of *yhiD* in *E. coli* did not impact ATP levels during cytoplasmic Mg^2+^ starvation (Fig. 2B). Taken together, these results indicate that most MgtC homologs function to control Pi and ATP homeostasis and that this is a conserved physiological function of these proteins.

### MgtC control of phosphate and ATP homeostasis during cytoplasmic Mg^2+^ starvation depends on genomic context

In contrast to the aforementioned results, we determined that deletion of *mgtC* in *S. praecaptivus* and *P. luminescens* did not hinder the growth of these species in media containing limiting Mg^2+^ (Fig. 3A and 3B). In agreement with this notion, wild-type and *mgtC* mutant strains of *S*. *praecaptivus* and *P. luminescens mgtC* exhibited equivalent ATP concentrations during growth in low Mg^2+^ media (Fig. 3C and 3D). Surprisingly, in both species, *mgtC* transcription was induced by Mg^2+^ starvation (Fig. 3E and 3F), indicating that these proteins still function under this stress, albeit not by altering Pi and ATP metabolism.

**Figure 3.**
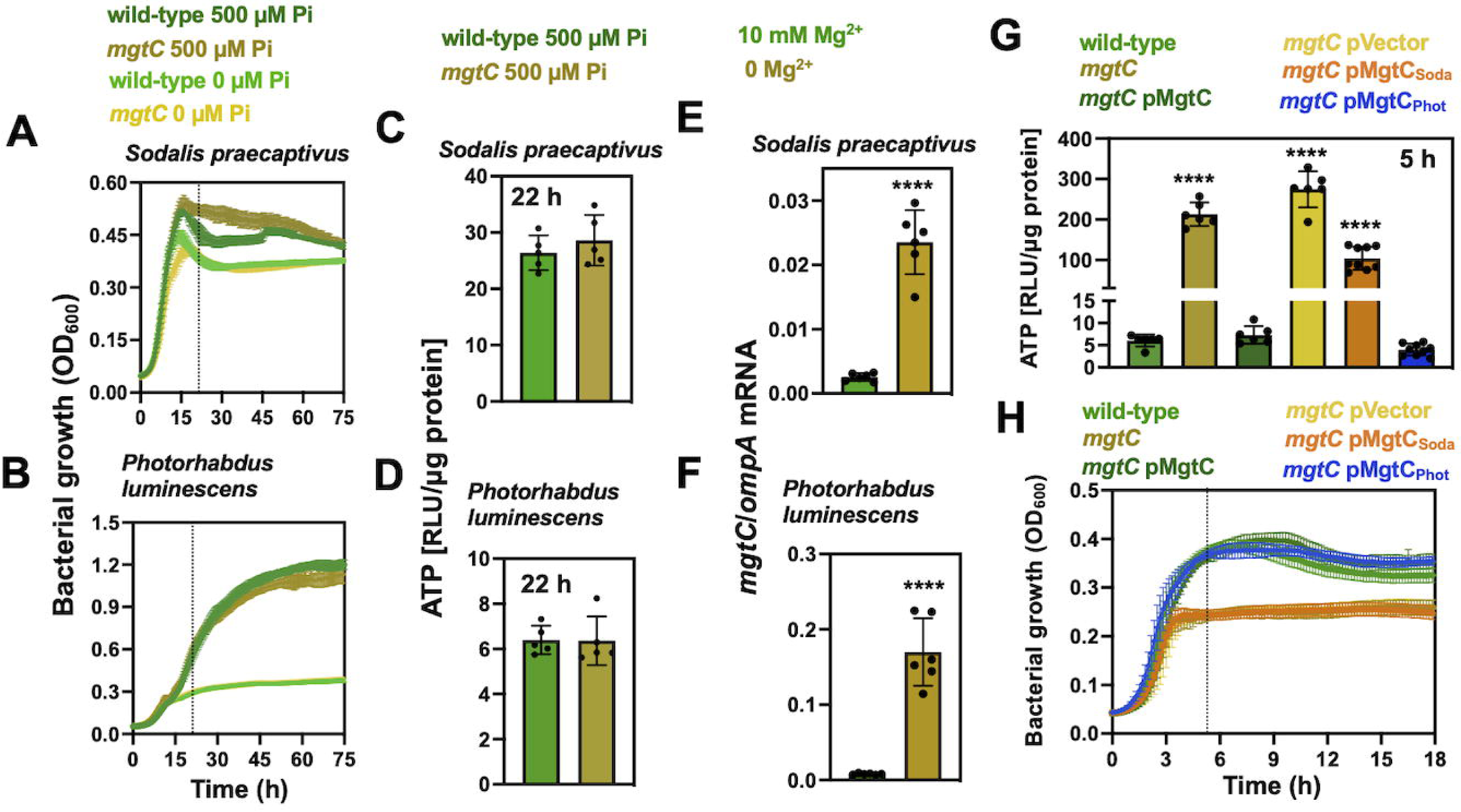
Effect of genomic and biochemical context on the function of MgtC homologs during magnesium starvation. Growth curves of **(A)** *S. praecaptivus* wild-type (ATCC BAA-2554) and isogenic *mgtC* mutant (RB61), and **(B)** *P. luminescens* wild-type (TT01) and isogenic *mgtC* mutant (RB101). Bacteria were grown in MOPS medium containing 10 µM MgCl_2_ and either 500 µM or no exogenous phosphate in the form of K_2_HPO_4_. Dashed lines indicate time points when intracellular ATP levels were measured. Relative intracellular ATP levels of **(C)** wild-type (ATCC BAA-2554) and isogenic *mgtC* mutant (RB61), and **(D)** *P. luminescens* wild-type (TT01) and isogenic *mgtC* mutant (RB101). ATP measurements were conducted in MOPS medium containing 10 µM MgCl_2_ and 500 µM of K_2_HPO_4_ at the indicated time points. *mgtC* mRNA levels in **(E)** *S. praecaptivus* (ATCC BAA-2554) and **(F)** *P. luminescens* (TT01) following 2 h incubation in MOPS medium containing 2 mM K_2_HPO_4_ and either 10 mM or no MgCl_2_. In both graphs, *mgtC* transcripts are normalized by transcripts from the housekeeping gene *ompA*. **(G)** ATP measurements of *S. enterica* wild-type (14028s), *mgtC* (EL4) and *mgtC* (EL4) strain harboring either a plasmid expressing *S. enterica*’s *mgtC* (pMgtC), *S. praecaptivus’*s *mgtC* (pMgtC_Soda_), *P. luminescens’ mgtC* (pMgtC_Phot_), or the empty vector pUHE-21 (pVector). Strains were grown in MOPS medium containing 10 µM MgCl_2_ and 500 µM of exogenous phosphate in the form of K_2_HPO_4_ as outlined in (H). (**H**) Growth of strains depicted in (G). Dashed line indicates the time point where intracellular ATP levels shown in (G) were measured. For (C-F): ****P < 0.0001, unpaired t-tests between sample populations. For (G): ****P < 0.0001, Two-way ANOVA calculated with Dunnett multiple-comparison test; statistical comparisons were made between wild-type *S. enterica* (14028s) and each of the genotypes. For all graphs means ± SDs of at least three biological replicates are shown.

Interestingly, several MgtC homologs, including that of *P. luminescens’*s, were shown to rescue the growth defect of an *S. enterica mgtC* mutant in low Mg^2+^ medium [30]. Consequently, we decided to test whether the MgtC from *S. praecaptivus* and *P. luminescens* could prevent the toxic accumulation of ATP in a *S. enterica mgtC* mutant. We established that the MgtC from *S. praecaptivus* conferred a partial reduction in ATP levels (Fig. 3G), which was insufficient to rescue growth of the *S. enterica mgtC* mutant strain (Fig. 3H). In contrast, expression of the MgtC from *P. luminescens* normalized ATP (Fig. 3G) and restored the growth (Fig. 3H) [30] of the *S. enterica mgtC* mutant to wild-type levels. These results demonstrate that the physiological role of MgtC is not intrinsic to the protein. While the physiological functions of some MgtC homologs have expectedly diverged, the roles these proteins play are conditioned by the cellular environment in which they operate.

### MgtC control of intramacrophage replication depends on genomic context

In *S. enterica*, MgtC promotes virulence by enabling intramacrophage replication [22, 51, 61]. While MgtC homologs can foster virulence in other species, these proteins do not always enhance intramacrophage replication [30, 35]. Considering the abovementioned results, we wondered if MgtC homologs that do not promote intramacrophage replication in their current resident species could confer this property to a *S. enterica mgtC* mutant. We evaluated the impact of the MgtC proteins from the closely related species *K. pneumoniae* and the distantly related species *B. thailandensis* on intramacrophage replication. We determined that the inactivation of *mgtC* from either *K. pneumoniae* or *B. thailandensis* did not affect the ability of these species to replicate in J774A.1 macrophage-like cells (Fig. 4A and B). This likely reflects specific physiological demands and pathogenic strategies that these species evolved to survive and replicate in their phagosome environments. Notably, heterologous expression of the *mgtC* from either of these species promoted intramacrophage replication of a *S. enterica mgtC* mutant (Fig. 4C). These results indicated that despite being dispensable for intramacrophage replication in their current species, the MgtC homologs from *K. pneumoniae* or *B. thailandensis* retain the genetic information required to promote intramacrophage replication in *S. enterica mgtC* mutant strains.

**Figure 4.**
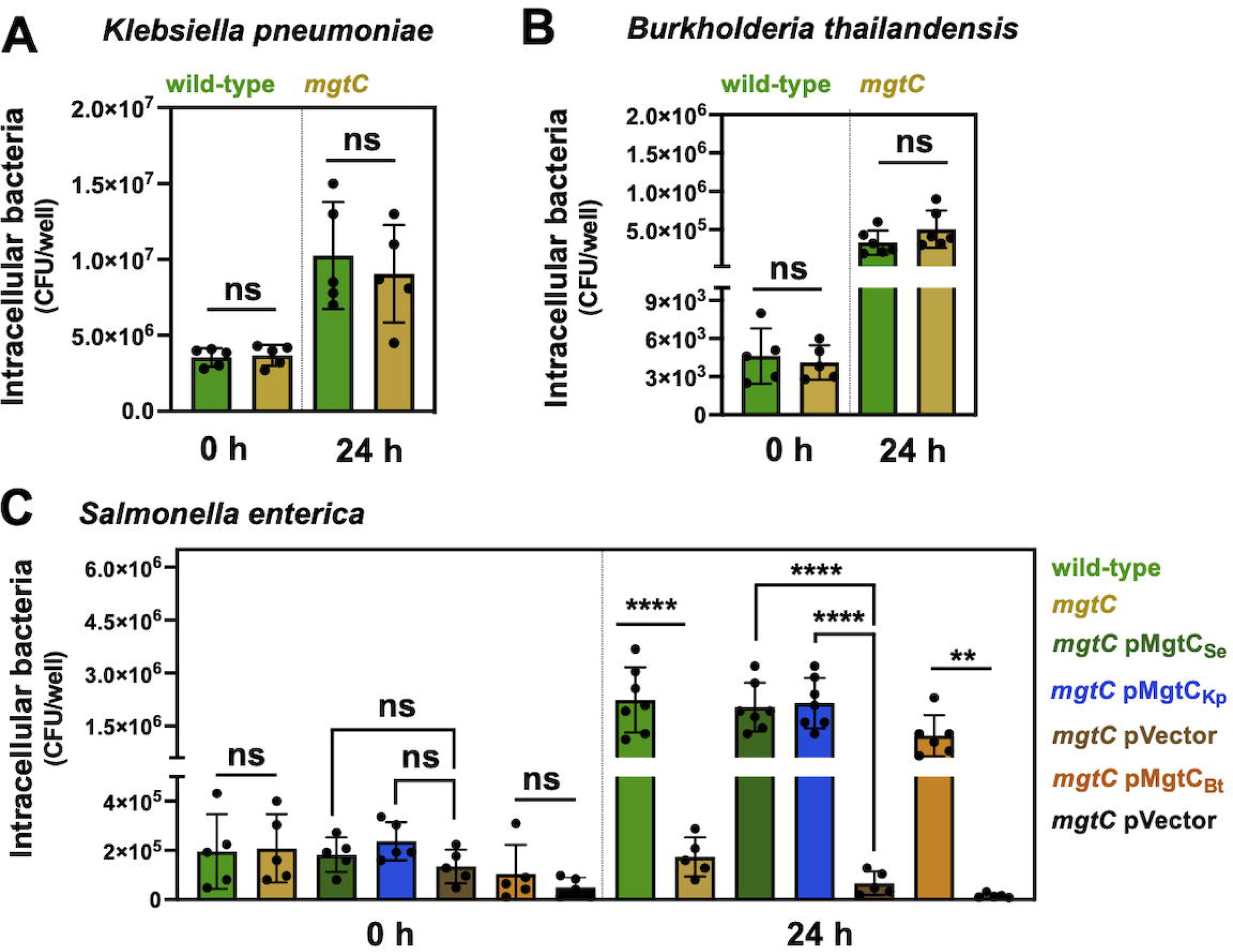
Effect of genomic context on the function of MgtC homologs during intramacrophage replication. Replication within J774A.1 macrophages of **(A)** wild-type (MKP103) and *mgtC* mutant (KP5811) strains of *Klebsiella pneumoniae*, and **(B)** wild-type (E264) and *mgtC* mutant (MP1936) strains of *Burkholderia thailandensis*. **(C)** Replication within J774A.1 macrophages of wild-type (10428s), *mgtC* mutant (EL4), and *mgtC* mutant (EL4) strain harboring plasmids expressing *mgtC* homolog from *Salmonella enterica* pMgtC (p18-MgtC_Se_), *K. pneumoniae* pMgtC_Kp_ (p18-MgtC_Kp_), *B. thailandensis* pMgtC_Bt_ (pBAD18-MgtC_Bt_), or empty vector controls pVector (p18 and pBAD18). ****P < 0.0001 and **P < 0.01, One-way ANOVA; P-values are shown for comparisons between wild-type and *mgtC*, *mgtC* pVector (p18 plasmid) and either *mgtC* pMgtC_Se_ (p18-MgtC from *S. enterica*) or *mgtC* pMgtC_Kp_ (p18-MgtC from *K. pneumoniae*), and *mgtC* pVector (pBAD18) and or *mgtC* pMgtC_Bt_ (pBAD18-MgtC from *B. thailandensis*) populations. Graphs depict means ± SDs of biological replicates. Results are representative of at least three independent experiments.

### The role of MgtC on *Salmonella enterica* Pi metabolism is context-dependent

Given the effect of genomic context on the role of MgtC homologs (Fig. 3 and 4), we wondered if we could identify context-dependent MgtC-regulated physiological processes in *S. enterica*. Because MgtC-inhibition of Pi metabolism is essential for survival to cytoplasmic Mg^2+^ starvation [53, 54, 62], but Pi metabolism has a mild fitness effect during infection [51, 63–65], we hypothesized that MgtC may not inhibit Pi metabolism inside macrophages. To address this possibility, we took advantage of the following phenomenon: in *S. enterica*, MgtC expression inhibits Pi uptake, thereby decreasing free cytoplasmic Pi and activating the PhoB/PhoR two-component signal transduction system. This phenomenon can be measured as green fluorescence emitted from a PhoB-activated *pstS*-*gfp* transcription reporter fusion (Fig. 5A) [62]. This behavior takes place when MgtC is expressed from its native promoter and chromosomal location in response to cytoplasmic Mg^2+^ starvation (Fig. 5A), but it can also be elicited by heterologous MgtC overexpression when cytoplasmic Mg^2+^ is not limited (Fig. 5B) [53, 54, 62].

**Figure 5.**
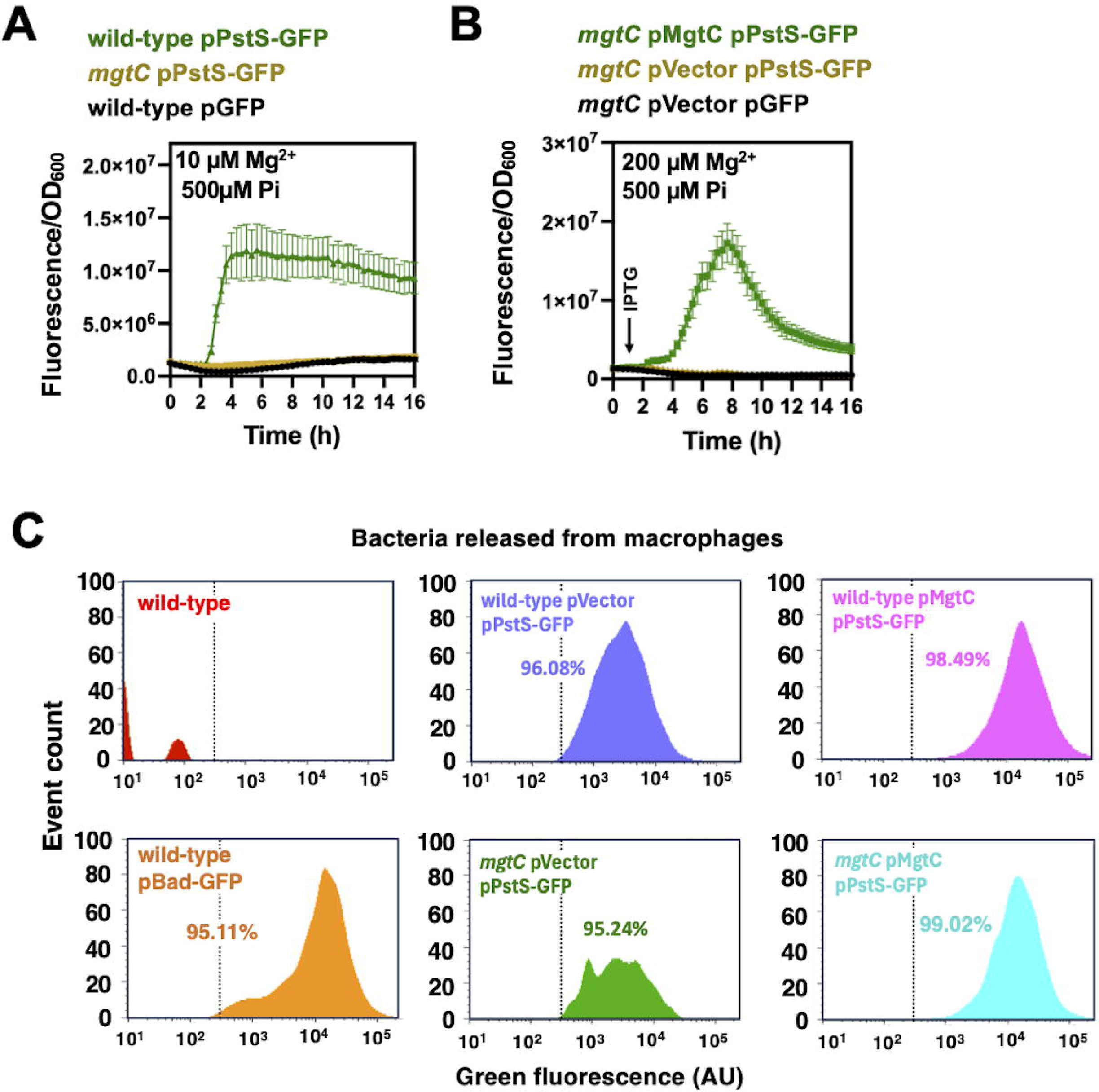
Effect of MgtC on Pi starvation response in *Salmonella enterica* replicating in macrophages. Effect of *mgtC* expression on the activity of a plasmid-borne *pstS-gfp* transcriptional reporter fusion. **(A)** Kinetic fluorescence measurements derived from *S. enterica* wild-type (14028s) and *mgtC* mutant (EL4) strains harboring pPstS-GFP (pACYC-P*pstS*-GFPc) or *S. enterica* wild-type (14028s) harboring a promotorless GFP plasmid (pGFP). Bacteria were grown in MOPS medium containing 10 µM MgCl_2_ and 500 µM of K_2_HPO_4_. **(B)** Kinetic fluorescence measurements derived from *S. enterica* wild-type (14028s) and *mgtC* mutant (EL4) strains harboring either pMgtC pPstS-GFP (pUHE-MgtC pACYC-P*pstS*-GFPc), pVector pPstS-GFP (pUHE-21 pACYC-P*pstS*-GFPc), or pVector pGFP (pUHE-21 pACYC-GFPc). **(C)** Fluorescent-activated cell sorting of mCherry-expressing populations of *S. enterica* released from J774A.1 macrophages. Graphs display green fluorescence of wild-type (SL233) or *mgtC* mutant (SL234) harboring either pVector pPstS-GFP (p18 pACYC-P*pstS*-GFPc) or pMgtC pPstS-GFP (p18-MgtC pACYC-P*pstS*-GFPc). The negative control, consisting of a population of non-fluorescent wild-type (14028s), and the positive control, consisting of a population of wild-type (14028s) expressing GFP from an arabinose-inducible plasmid (pBAD18-GFP), are also shown. The percentage of GFP^+^ bacteria within each population is displayed. Results are representative of at least three independent experiments. Graphs show means ± SDs of at least three biological replicates.

Accordingly, we compared the fluorescence emitted by a *pstS-gfp* reporter in the wild-type and *mgtC* null mutant strain, each carrying either a multicopy plasmid overexpressing MgtC (pMgtC) or an empty vector (pVector) control. We determined that the distribution and intensity of fluorescence were indistinguishable between wild-type pVector pPstS-GFP and *mgtC* pVector pPstS-GFP populations (Fig. 5C), indicating that expression of MgtC from its native chromosomal location does not affect *pstS-gfp* activity, as observed when bacteria experience cytoplasmic Mg^2+^ starvation in laboratory medium (Fig. 5A). In contrast, plasmid overexpression of MgtC in populations of wild-type pMgtC pPstS-GFP and *mgtC* pMgtC pPstS-GFP increased the fluorescence intensity of cells in these populations, with negligible impact on the distribution of fluorescent bacteria (Fig. 5C). As expected, fluorescence was negligible in a non-fluorescent strain (wild-type) and elevated in a strain expressing GFP from a plasmid-borne arabinose-inducible (wild-type pBAD-GFP) (Fig. 5C). Together, this data establishes that while MgtC inhibits Pi metabolism during growth in defined laboratory medium, it does not perform this function when *S. enterica* replicates in macrophages.

### The role of MgtC on *Salmonella enterica* translation homeostasis is dependent on the biochemical context

When *S. enterica* experiences cytoplasmic Mg^2+^ starvation during growth in defined laboratory media, MgtC promotes a decrease in Pi metabolism and ATP production, which is required for normal translation. Consequently, an *mgtC* mutant is unable to inhibit Pi metabolism and ATP synthesis, displaying ribosomal subunit assembly defects and a decreased translation rate [52–54] (Fig. 6A). This decreased translation rate can be reversed by artificially hydrolyzing ATP or hindering the access of bacteria to Pi. This implies that decreased translation observed in this strain results from elevated Pi assimilation [52–54].

**Figure 6.**
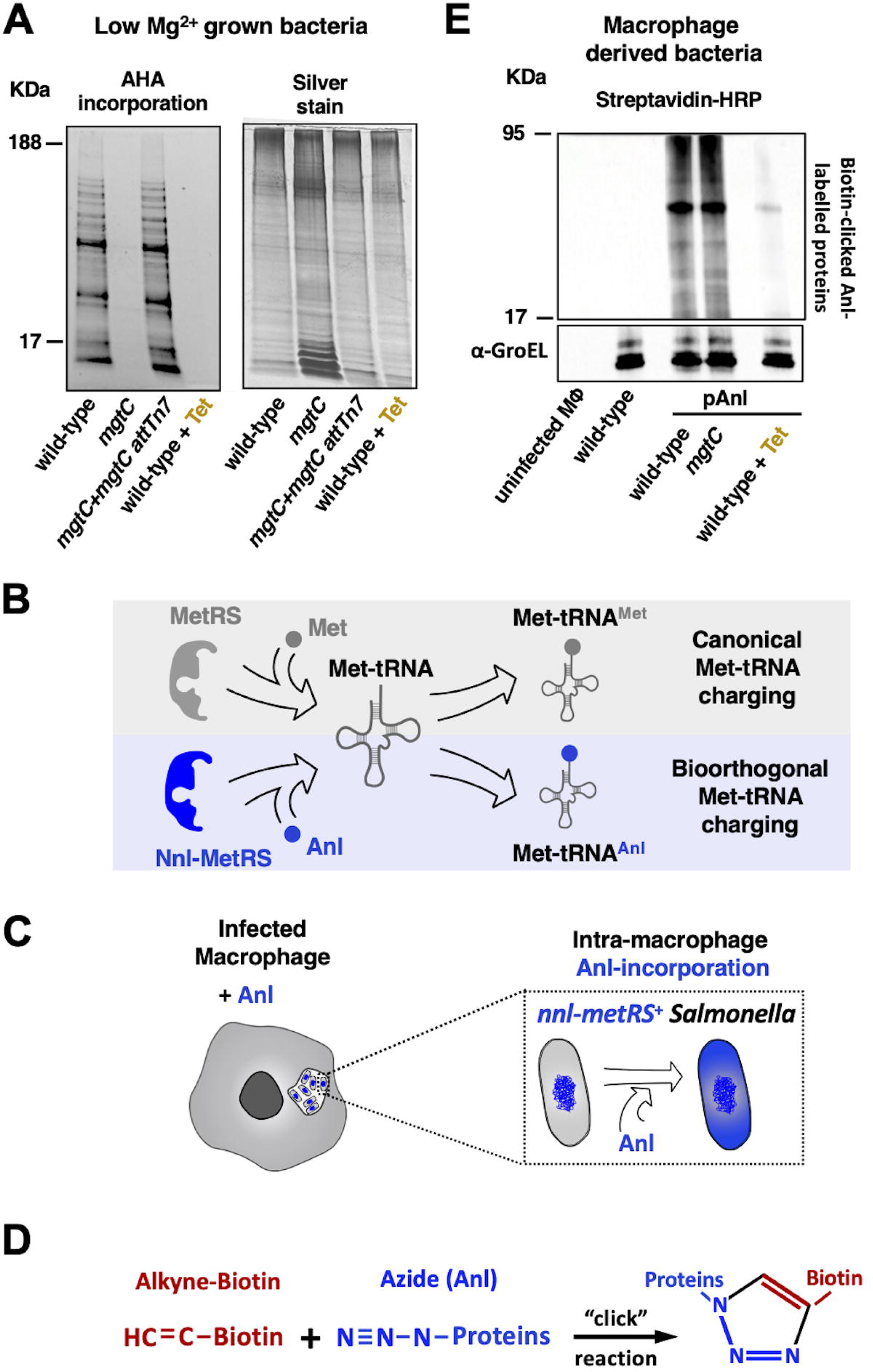
Effect of MgtC on *Salmonella enterica* translation during replication in macrophages. **(A)** Resolved sodium dodecyl sulfate–polyacrylamide gel containing L-azidohomoalanine (AHA) labeled protein samples derived from *S. enterica* wild-type (14028s), *mgtC* (EL4), and a complemented *mgtC* mutant expressing *mgtC* from the selectively neutral Tn*7* attachment site (*mgtC+mgtC* attTn*7,* RB391). Cells were grown for 5 h in MOPS medium containing a defined amino acid mixture lacking methionine (see Supplemental Material), 10 µM MgCl_2_ and 2 mM of K_2_HPO_4_, prior to being labeled with AHA. The left-hand side depicts AHA-labeled proteins clicked with TAMRA-alkyne. The right-hand side depicts the same gel following silver staining. **(B-D)** Cartoons depicting the strategy used to measure protein synthesis of intracellular bacteria. **(B)** The methionyl-tRNA synthetase Nnl-MetRS catalyzes the bioorthogonal charging of Met-tRNA with the non-canonical clickable amino acid L-azidonorleucine (Anl), thereby enabling the incorporation of Anl into proteins. **(C)** When macrophages are infected with Nnl-MetRS-expressing *S. enterica* strains, the addition of Anl to the culture medium allows intracellular bacteria to incorporate this amino acid into newly synthesized proteins. **(D)** Following Anl labeling, Anl-containing proteins in cell lysates can be covalently linked to an alkyne-containing detection group (e.g. Alkyne-biotin) through a copper-catalyzed azide–alkyne cycloaddition. **(E)** Western blots of protein samples derived from J774A.1 macrophages infected with either wild-type (14028s), *mgtC* (EL4) harboring pAnl. Bacteria were allowed to propagate for 12 h, then labeled for 12 h with the addition of 1 mM Anl to the macrophage medium. Control samples consisting of uninfected macrophages, macrophages infected with wild-type (14028s) lacking plasmid pAnl, or wild-type pAnl and incubated with Anl in the presence of 100 µg/ml of tetracycline are also shown. Total protein samples were clicked to biotin-alkyne and analyzed by western blot. The top portion of the membrane depicts total biotin-clicked proteins probed with streptavidin conjugated to Horseradish Peroxidase (streptavidin-HRP). The same membrane was stripped and probed with antibodies against *S. enterica* GroEL (*a*-GroEL). Signals derived from *a*-GroEL are used as proxies for bacterial protein in each sample. Gels and membranes are representative of at least 3 independent experiments.

Because MgtC does not affect Pi metabolism during replication in macrophages (Fig. 5C), we reasoned that the wild-type and *mgtC* mutant strains of *S. enterica* would display similar translation rates inside these cells. To test this idea, we engineered a series of *S. enterica* strains with an expanded genetic code [66]. These strains harbored the pAnl plasmid, which expresses a methionyl-tRNA synthetase (*nnl-metRS*) capable of biorthogonal incorporation of the unnatural clickable amino acid L-azidonorleucine (Anl) into nascent proteins (Fig. 6B). Addition of Anl to the cultures of macrophages infected with pAnl harboring *S. enterica* enable selective incorporation of this amino acid by intracellular bacteria (Fig. 6C), and subsequent detection via the introduction of a detection group through a cooper catalyzed click-reaction (Fig. 6D).

We established that a pAnl-harboring wild-type and *mgtC* mutant strains of *S. enterica* incorporated similar amounts of Anl into proteins when replicating in macrophages grown in medium supplemented with Anl (Fig. 6E). As expected, uninfected macrophages or macrophages infected with wild-type *S. enterica* lacking the pAnl plasmid were unable to incorporate Anl into proteins (Fig. 6E). Importantly, the addition of the bacterial translation inhibitor tetracycline to the macrophage medium drastically decreased Anl incorporation (Fig. 6E); thus demonstrating that Anl was incorporated by the ribosomes of intracellular *S. enterica*. Taken together, these results establish that while MgtC promotes translation homeostasis when *S. enterica* is grown in defined culture medium containing limiting Mg^2+^ it does not affect translation during growth inside macrophages.

## Discussion

In the current study, we demonstrate that the roles of the MgtC protein of *S. enterica* in regulating Pi and ATP metabolism during cytoplasmic Mg^2+^ starvation are broadly conserved. We show that while some MgtC homologs have divergent roles in certain species, these proteins can assume this function when expressed in *S. enterica*. We establish that this phenomenon also occurs during intramacrophage replication. Specifically, whereas some MgtC homologs do not contribute to intramacrophage replication in their current species, they can confer this property to *S. enterica*. Finally, we show that MgtC controls Pi homeostasis and translation when *S. enterica* cells experience cytoplasmic Mg^2+^ starvation in a defined culture medium, but not when bacteria reside inside macrophages.

The MgtC protein of *S. enterica* was initially shown to promote survival to Mg^2+^ starvation in laboratory medium and enable replication in mammalian macrophages [22, 45]. While these phenotypes were largely recapitulated in other bacterial species [22, 26, 28, 30, 33–36, 40–43, 45],

*S. enterica*’s MgtC was eventually shown to be expressed in response to intracellular signals that are triggered by a decrease in cytoplasmic Mg^2+^ [47–50, 59, 67, 68], and to reduce ATP concentrations by physically interacting with the AtpB protein component of *S. enterica*’s FoF1 ATP synthase [24]. More recently, MgtC was also shown to interact with *the* high-affinity Pi transporter complex of *S. enterica*, PstSCAB-PhoU-PhoR, and to stimulate transport during Mg^2+^ starvation [69]. However, while MgtC may interact with components of this Pi transporter, the outcome of these interactions must contend with the facts that excessive Pi leads to ATP overproduction, which is acutely toxic to bacteria experiencing cytoplasmic Mg^2+^ starvation, and that *mgtC*-encoding bacterial species often employ this gene product to alleviate Pi-induced ATP toxicity [53, 54] (Fig. 2). Additionally, our results indicate that ectopic plasmid over-expression of MgtC [69] creates non-physiological interactions and artificial PhoB activation (Fig. 5).

In *S. enterica,* the genetic inactivation of *mgtC* results in several phenotypes, including ATP accumulation, when bacteria experience cytoplasmic Mg^2+^ starvation during growth in laboratory media [22, 25, 30, 45, 52–54]. Because the artificial normalization of ATP levels in an *mgtC* null background reverts many of these phenotypes, MgtC has been proposed to function in ATP homeostasis [24]. Nonetheless, the results presented herein support the notion that the pleiotropic effects of MgtC arise from multiple regulatory interactions that alter the activities of distinct proteins and the physiological processes they regulate. Accordingly, MgtC interactions that are posited to occur when *S. enterica* experiences cytoplasmic Mg^2+^ starvation in laboratory medium enable this bacterium to inhibit Pi metabolism, prevent ATP accumulation, and maintain translation homeostasis. However, MgtC does not affect Pi metabolism (Fig. 5) or translation during intramacrophage replication (Fig. 6). This likely reflects that the physiological burdens countered by MgtC during cytoplasmic Mg^2+^ starvation in laboratory medium differ from those it mitigates in the macrophage phagosome.

Consistent with this notion, MgtC is required for *S. enterica* replication in macrophages that lack the Nramp1 (Slc11a1) gene, which encodes a divalent transition metal transporter that is expressed on the membranes of the *<u>S</u>almonella*-<u>c</u>ontaining <u>v</u>acuole (SCV), and that restricts Mg^2+^ availability within this compartment [31, 60, 70]. This indicates that MgtC does not respond to cytoplasmic Mg^2+^ starvation during replication in macrophages. Furthermore, *S. enterica* mutants defective in either the Pi starvation response or purine biosynthesis are attenuated in intramacrophage replication [61, 71]. This suggests that the metabolic precursors of ATP are scarce in the phagosome, and that *S. enterica* is unlikely to experience ATP intoxication and subsequent disruption of ribosome assembly inside these phagocytic cells—a notion that is supported by our experimental results (Fig. 5C and Fig. 6). While this indicates that MgtC is a regulator, it also implies that the regulatory interactions governing Pi and translation homeostasis that take place in low Mg^2+^ laboratory medium either do not occur, or do not impact bacterial fitness in the macrophage phagosome [30] (Fig. 5 and 6).

The MgtC homologs of *B. thailandensis* and *K. pneumoniae* are dispensable for intramacrophage replication in these species. This likely reflects that these bacteria occupy different intramacrophage environments and have evolved distinct mechanisms to survive and replicate within these cells, rendering MgtC’s function redundant in these physiological contexts. Like *S. enterica*, *K. pneumoniae* can persist within specialized macrophage vacuoles by avoiding fusion with lysosomes and evading other bactericidal mechanisms deployed by the host cell [72–75]. While some of the conditions within the *Klebsiella*-containing vacuoles may resemble those found in the SCV, *K. pneumoniae* is primarily an extracellular pathogen and, unlike *S. enterica,* lacks an extensive repertoire of virulence factors that support efficient intramacrophage replication [76–79]. Consistent with the notion that MgtC is not a virulence factor in *K. pneumoniae*, its transcription is not induced during mouse infection [80].

Compared to *K. pneumoniae* and *S. enterica*, *B. thailandensis* does not remain in a phagosome. Shortly following internalization, this bacterium lyses the macrophage vacuole and escapes into the nutrient-rich cytosol to replicate [81–83]. Accordingly, *B. thailandensis* does not express *mgtC* in macrophages [84], and its deletion does not impair this bacterium’s ability to replicate in these host cells (Fig. 4B). This contrasts with the behavior of the closely related *B. cenocepacia*, which largely remains within a macrophage phagosome and requires *mgtC* for intracellular replication [33, 85, 86].

While the MgtC homologs from either *B. thailandensis* or *K. pneumoniae* do not impact their replication within murine macrophages, these proteins can promote the replication of a *S. enterica mgtC* null mutant in these phagocytic cells (Fig. 4C). That is, they contain the genetic information that is also present within *S. enterica*’s native MgtC and that is required for this bacterium to replicate in its intramacrophage environment. Similarly, the MgtC from *P. luminescence* does not affect ATP homeostasis or the growth of this bacterium in low Mg^2+^ laboratory medium. Nonetheless, this protein retains the information that is required to regulate these processes in *S. enterica* (Fig. 3) [30]. These results suggest that the physiological roles of MgtC arise from regulatory interactions with cellular components present in *S. enterica*, that are absent in these other species.

Genetic diversification can promote the emergence of novel regulatory interactions. In bacteria, this often occurs through the horizontal acquisition of new genes [11–15, 19, 65, 87–91]. The fate of horizontally acquired genes is largely determined by the fitness effects that they confer on recipient cells. Natural selection typically purges deleterious genes and drives beneficial ones to fixation. Whereas selectively neutral genes are more susceptible to stochastic forces such as drift, in bacteria, redundant sequences are eventually eliminated by a mutational bias towards deletions [92, 93]. Taking this into consideration, it is tempting to speculate that recently acquired *mgtC* homologs may be incorporated into bacterial lineages by establishing regulatory interactions with the cellular components governing ancestrally controlled physiological processes, given that these components are likely to have inherited the information required for these interactions—for example, interacting sequence spaces within co-expressed proteins that regulate Pi uptake, ATP synthesis and other processes that promote survival to cytoplasmic Mg^2+^ starvation (Fig. 1-4). This may enable such lineages to radiate to previously inaccessible niches [1–5], where the integrity of these critical physiological processes could be protected by these newly established MgtC interactions. The subsequent formation of additional species-specific regulatory interactions could further alter the ancestral biochemical environments, facilitating the evolution of new traits and potentiating additional radiation events [1, 2].

## Materials and Methods

### Bacterial strains, plasmids, oligonucleotides and growth conditions

Bacterial strains and plasmids used in this study are listed in Table S1. Oligonucleotide sequences are presented in Table S2. Unless otherwise stated, routine growth and recombination experiments were carried out in following media: Luria-Bertani medium (*Acinetobacter baumannii*, *Escherichia coli*, *Klebsiella pneumoniae, Photorhabdus luminescens*, *Salmonella bongori*, *Salmonella enterica*, *Yersinia enterocolitica*), Luria–Bertani lacking exogenous sodium chloride (LSLB) medium (*Burkholderia thailandensis, Burkholderia cenocepacia* and *Pseudomonas aeruginosa, Sodalis praecaptivus*) or brain-heart infusion (BHI) medium (*Yersinia pestis*). When required, marked mutations in the *Salmonella enterica* chromosome were introduced into different genetic backgrounds via bacteriophage P22 transduction, as described [94]. Following genetic modification, bacterial clones were recovered by isolating single colonies on 1.5% (w/v) agar plates. *Photorhabdus luminescens, S. praecaptivus,* and *Y. pestis* were propagated at 30°C.

*Acinetobacter baumannii*, *B. cenocepacia, B. thailandensis*, *E. coli, K. pneumoniae*, *P. aeruginosa*, *S. bongori*, *S. enterica*, and *Y. enterocolitica* were propagated at either 30 or 37°C. When required, media was supplemented with the following: 200 μg/ml (*A. baumannii*) or 100 μg/ml of ampicillin, 30 μg/ml of apramycin, 20 μg/ml of chloramphenicol, 50 μg/ml (*E. coli*, *S. bongori*, *S. praecaptivus, Y. enterocolitica* and *Y. pestis*) or 35 μg/ml (*P. luminescens*) of kanamycin, 20 μg/ml (*E. coli* and *S. enterica*) or 70 μg/ml (*B. thailandensis* and *P. aeruginosa*) of tetracycline, 100 μg/ml of spectinomycin, 50 μg/ml of trimethoprim, 1 μg/ml of anhydrotetracycline, 60 μg/ml of diaminopimelic acid, 40 μM of L-azidohomoalanine (AHA), 2 mM (*A. baumannii*) or 250 μM of isopropyl β-d-1-thiogalactopyranoside (IPTG), 1 mM of cyclic adenosine monophosphate, and 0.1% (w/v) of L-arabinose.

### Construction of mgtC mutant strains of Salmonella bongori, Sodalis praecaptivus, Yersinia enterocolitica and Yersinia pestis

*Salmonella bongori*, *S. praecaptivus*, *Y. enterocolitica* and *Y. pestis ΔmgtC::km^R^* mutants were constructed by bacteriophage λ recombineering and purified *ΔmgtC::km^R^* PCR products [95, 96]. Detailed protocols for strain construction are outlined in the Supplemental Material.

### Construction of *mgtC* mutant strain of *Photorhabdus luminescens*

*Photorhabdus luminescens ΔmgtC::km^R^* was constructed through biparental mating with an *E. coli* strain harboring a *ΔmgtC::km^R^* allele in a transferable rep_RK2_ suicide plasmid. We followed the conjugation protocol described in [97] with modifications outlined in the Supplemental Material.

### Construction of *mgtC* mutant strain of *Acinetobacter baumannii*

*Acinetobacter baumannii ΔmgtC* was constructed using the *recBA* system and a purified *ΔmgtC::Apr* PCR product as described [98]. A detailed protocol for constructing this strain is provided in the Supplemental Material.

### Construction of *mgtC* mutant strain of *Burkholderia thailandensis*

*Burkholderia thailandensis ΔmgtC::*Tn*10* was constructed through natural competence [99]) using a purified *ΔmgtC::*Tn*10* PCR product. A detailed protocol for the construction of this strain is outlined in the Supplemental Material.

### Construction of plasmids

Plasmids were constructed using NEBuilder HiFi DNA Assembly (New England Biolabs), as described in the Supplemental Material.

### Construction of *Salmonella enterica* strains harboring insertions at the Tn*7* attachment site

Inserts from plasmids pGRG36-P*mgtC-leader-mgtC-*FLAG, pTn7-mCherry-P1 were moved into the Tn*7* attachment site in the *Salmonella enterica* chromosome using the standard protocol [100] with the modifications described in the Supplemental Material.

### Measuring bacterial growth

Bacterial growth was measured in MOPS-based media [101] in 96-well plates, in a SpectraMax i3x (Molecular Devices) plate reader as described in [62] with the modifications described in the Supplemental Material.

### Measuring ATP levels in bacterial samples

ATP measurements were carried out as described [25].

### mRNA quantification

Bacterial total RNA was extracted using the RNeasy Kit (Qiagen). cDNA was synthesized from RNA samples using the SuperScript VILO cDNA synthesis kit (Thermo Fisher Scientific) and quantified using a standard curve method in a QuantStudio 7 Flex Real-Time PCR detection system (Thermo Fisher Scientific). A detailed mRNA quantification protocol is described in the Supplemental Material.

### Measuring bacterial fluorescence during growth in defined media

*Salmonella enterica* strains were grown at 37°C and 250 rpms in MOPS medium containing 10 mM MgCl_2_ and 2 mM K_2_HPO_4_. Stationary-phase cultures were washed 3x with MOPS salts lacking magnesium and phosphate, then diluted 1:100 into fresh MOPS containing 200 μM MgCl_2_ and 500 μM K_2_HPO_4_. Bacterial cultures were aliquoted in black, clear-bottom, 96-well plates (Corning) and individual wells were sealed with two drops of mineral oil. Plates were propagated at 37°C with auto-mixing in a SpectraMax i3x (Molecular Devices) plate reader. Following 1 h of growth, cultures were treated with 250 µM IPTG. Green fluorescence (excitation 485 nm/emission 535 nm) and absorbance at 600 nm (OD600) were measured for each well at regular time intervals. Fluorescence values were normalized by the OD600 of the samples from each well.

### Measuring bacterial translation rates during growth in defined media

Translation rates in bacteria grown in MOPS medium were determined by measuring the incorporation of the non-canonical, clickable amino acid L-azidohomoalanine (AHA) into newly synthesized proteins as described previously [52]. Briefly, *Salmonella enterica* strains were grown overnight in MOPS medium [101], lacking CaCl_2_, and containing high (10 mM) MgCl_2_, 22 mM glucose and 0.1% casa amino acids. Overnight cultures were washed 3x in MOPS medium salts, and inoculated 1:100 into fresh medium lacking CaCl_2_ and containing low (10 μM) MgCl_2_, 22 mM glucose and an amino acid mixture consisting of 3.2 mM of alanine, glycine, leucine, glutamate and serine; 2.4 mM glutamine, isoleucine and valine; 1.6 mM arginine, asparagine, aspartate, lysine, phenylalanine, proline, and threonine; 0.8 mM histidine and tyrosine, and 0.4 mM cysteine and tryptophan. Cells were grown for 5 h at 37°C and 250 rpms and subsequently labeled for 30 min with 40 µM of AHA (Click Chemistry Tools). Click reactions were carried out, quantified and visualized as described [52].

### Intramacrophage replication assay

J774a.1 mouse macrophages were propagated in DMEM medium supplemented with 10% fetal bovine serum (FBS) at 37°C and 5% CO_2_. For macrophage invasion assays, these cells were seeded into 6-well plates (at a 1.5 x 10^6^ cells/well), 12-well plates (at a 1.5 x 10^5^ cells/well) or 24-well plates (at 5 x 10^5^ cells/well). After estimating the number of macrophage cells/well using a Cytosmart Cell Counter (Corning), macrophages were infected with bacteria at an MOI of 1 or 10 using a standard invasion assay. Briefly, macrophage medium was replaced with medium containing the desired number of bacteria, plates were centrifuged for 10 min at 1,000 rpm and then incubated at 37°C and 5% CO_2_ for 1 h to allow bacterial internalization. Macrophage monolayers were then washed three times with phosphate-buffered saline (PBS), and the remaining extracellular bacteria were killed by incubating the monolayers for 1 h in medium containing 100 μg/mL of gentamycin (*S. enterica* and *K. pneum*oniae) or 600 μg/mL of kanamycin (*B. thailandensi*s). Monolayers were then washed three times with PBS. For a subset of wells, macrophages were lysed by incubating the monolayer in PBS containing 1% Triton X-100 for 10 min and subsequently mixing by pipetting multiple times. Macrophage lysates were then diluted and plated to estimate the number of bacteria internalized at time 0. Macrophages in the remaining wells were incubated for 20 h in DMEM medium supplemented with 10% FBS, and either 12.5 μg/mL of gentamycin (*K. pneumoniae* and *S. enterica*) [25] or 250 μg/mL of kanamycin [102], at 37°C and 5% CO_2_. Monolayers were then washed twice with PBS and lysed with PBS 1% Triton X-100 as described above. The number of intracellular bacteria at 20 h post-infection was estimated by plating lysate dilutions as described above.

### Measuring bacterial fluorescence during growth in macrophages

J774A.1 macrophages were seeded in 6-well plates and infected with bacteria at an MOI of 20 for 30 min. Following infection, cells were washed three times with PBS and incubated in DMEM supplemented with 10% FBS and 100 µg/mL gentamicin for 1 h to eliminate extracellular bacteria. The medium was then replaced with DMEM containing 10 µg/mL gentamicin for the remainder of the experiment. To induce GFP expression cloned into plasmid pBAD18 [103], 0.1% L-arabinose was added to the macrophage medium. At 18 h post-infection, macrophages were washed twice with PBS, and intracellular bacteria were released by lysis in PBS containing 1% Triton X-100. Cellular debris was removed by centrifugation at 1000 × g for 3 min, and bacteria in the supernatant were collected by centrifugation at 14,000 × g for 1 min. Pellets were washed three times with PBS, gently resuspended in 0.5 mL PBS, and fixed in 0.5 mL ice-cold 4% paraformaldehyde for 15 min at room temperature. Fixed bacteria were washed twice and resuspended in FACS buffer (PBS with 1% FBS and 1 mM EDTA). Samples were stored at 4 °C, protected from light, and analyzed within 24 h. Flow cytometry was performed on a BD LSRFortessa (BD Biosciences, San Jose, Calif.), Penn State College of Medicine’s Flow Cytometry Core (RRID:SCR_021134). GFP was detected in the FITC channel and mCherry in the PE-Texas Red-A channel. Compensation was performed using single-fluorescent protein bacterial controls. At least 100,000 events were acquired per sample. Data were analyzed using FlowJo v11.1. Bacteria were first gated on FSC/SSC to exclude debris, followed by singlet discrimination. The mCherry-positive gate was defined using the non-fluorescent wild-type parental strain as a negative control. GFP expression was quantified within the mCherry-positive bacterial population.

### Measuring bacterial translation rates during replication in macrophages

J774A.1 macrophages were infected with bacteria at an MOI of 100. At 12 h post-infection, samples were treated with 1 mM I-azidonorleucine (Anl). When needed, 100 μg/ml of tetracycline was added one h before Anl labeling. After 12 h of incubation at 37°C and 5% CO_2_, samples were washed twice with PBS and lysed with PBS containing 1% triton X-100 as described above. Samples were vortexed and bacteria were collected by centrifuging the samples for 5 min at 18,000 x g, and bacteria were resuspended in PBS. A fraction of the samples was diluted to estimate bacterial counts by colony-forming unit (CFU) counts. The remaining of the samples were collected by centrifugation for 5 min at 18,000 x g, and stored at -80°C. Frozen bacterial pellets were resuspended in lysis buffer (50 mM Tris pH 8.0, 1% sodium dodecyl sulfate (SDS)), and equal numbers of bacterial cells were disrupted by sonicating the samples in an ice bath. Cell debris were removed by centrifuging the samples for 5 min at 4°C and 18,000 x g. Proteins in the supernatant were precipitated with methanol-chloroform following the protocol described in the Click-iT Protein Reaction Buffer Kit (Thermo Fisher Scientific). Samples were resuspended in 50 mM Tris pH8 and 0.1% SDS, Protein concentrations were estimated using the Rapid Gold BCA Protein Assay kit (Thermo Fisher Scientific). Equal amounts of proteins were used in click-reactions with a biotin-Alkyne using the Click-iT Protein Reaction Buffer Kit (Thermo Fisher Scientific). Samples were resuspended and normalized in running buffer (50 mM Tris pH8, 1% SDS), mixed with SDS-PAGE loading buffer, heat-denatured and resolved in 4-12% Bis-Tri<u>s</u> gels.

Protein samples were transferred to a nitrocellulose membrane using an iBlot 3 transfer system (Invitrogen). The membrane was blocked for 1 h at room temperature with BSA blocking buffer (1% bovine serum albumin (BSA) fraction V, 0.2% (w/v) triton X-100 in PBS), probed for 1 h, at room temperature, with a streptavidin-HRP conjugated antibody (Sigma Millipore) diluted in BSA blocking buffer, and washed 6 times for 10 min with ABS blocking buffer (10% adult bovine serum, 1% (w/v) triton X-100 in PBS). Membrane-bound signals derived from HRP were detected with SuperSignal ELISA Femto Kit (Thermo Fisher Scientific) and captured with LAS-4000 (GE Healthcare) or an Azure 600 (Azure) imager. The membrane was subsequently stripped by incubation in 20 mL of stripping buffer (20 mL 10% SDS, 12.5 mL 0.5 M Tris-HCl, pH 6.8, 0.8 mL β-mercaptoethanol, and distilled water to a final volume of 100 mL) at 50°C for 15 min with gentle agitation, washed extensively with distilled water for 15 min, followed by a 5-min wash with PBST, and blocked with 5% (w/v) skim milk in PBST for 1 h at room temperature. The membrane was subsequently probed with an anti-GroEL primary antibody (Abcam) and HRP-conjugated anti-rabbit IgG secondary antibody (Cytiva). GroEL signal was detected as described above. The amount of GroEL in each sample was used as a loading control for bacterial cell lysates.

### Phylogenetic reconstruction

16S rDNA, *mgtC* and *yhiD* sequences were obtained from whole or partially assembled genomes of selected bacterial species that were deposited in the GenBank database. Accession numbers for these nucleotide sequences are listed in the Supplemental Material. 16S rRNA nucleotide and *mgtC* /*yhiD* codon sequences were aligned in the Molecular Evolutionary Genetics Analysis (MEGA) software using the ClustalW tool. Alignments were used to infer an optimal model of sequence evolution. Sequence distances were calculated using the Tamura 3-parameter model, with gamma-distributed rates and invariable sites. Initial trees were generated using Maximum Parsimony, and Maximum Likelihood trees were calculated using the Nearest-Neighbor-Interchange heuristic method with 10,000 bootstrap replicates.

### Statistical Analyses

Results were obtained from at least three independent experiments and were plotted using GraphPad PRISM. Data were analyzed in GraphPad PRISM using either an unpaired t-test, one-way or two-way ANOVA as indicated in each figure legend.

### Image Acquisition, Analysis, and Manipulation

Gel and membrane images were acquired using either an Amersham Imager 600 (GE Healthcare Life Sciences) or an Azure 600 Imager (Azure). ImageJ [104] was used to crop the images and adjust their brightness and contrast. These modifications were simultaneously performed across the entire set images to be shown.

## Supplemental Material

## Acknowledgements

We are grateful to Miguel A. Valvano (Queen’s University Belfast) for providing the wild-type and *mgtC* mutant strains of *Burkholderia cenocepacia*, and Virginia Miller (University of North Carolina) for providing a wild-type strain of *Yersinia enterocolitica*, and Bryan W. Davies (University of Texas at Austin) for providing plasmids pAT02 and pAT03. The authors are also grateful to two anonymous reviewers for their insightful comments and useful suggestions.

## Funding and additional information

M.H.P. is partially supported by Grant AI148774 from the NIH and funds from The Pennsylvania State University College of Medicine.

