## Supplemental Material for "Genomic and biochemical contexts determine the physiological role of a horizontally acquired gene"

### Tables

**Table S1. Bacterial strains and plasmids used in this study**

| Strain | Relevant characteristics | Identifier | Source |
|---|---|---|---|
| ***Escherichia coli*** | | | |
| EC100D | λ$pir^+$ (DHFR) host strain used for generation and propagation of plasmid constructs. | N/A | Epicentre |
| NEB5α | Host strain used for cloning. | N/A | New England Biolabs |
| BW29427 | Donor strain in matings, F-, RP4-2($Tet^S$, kan1360::FRT)?, *thrB1004*?, *ΔlacZ58*(M15)?, Δ*dapA1341*::[erm λ $pir^+$]?, *rpsL*($str^R$)?, *thi*-?, *hsdS*-?, *pro*-? | N/A | Yale University *Escherichia coli* Genetic Stock Center |
| MG1655 | wild-type | wild-type | (Blattner et al., 1997) |
| MP751 | *ΔyhiD::*$km^R$ | *yhiD* | (Pontes et al., 2016) |
| ***Acinetobacter baumannii*** | | | |
| ATCC 17978 | wild-type | wild-type | ATCC |
| A1S_2069 | *ΔmgtC* | *mgtC* | This study |
| ***Burkholderia cenocepacia*** | | | |
| K56-2 | wild-type | wild-type | (Maloney & Valvano, 2006) |
| KEM1 | *mgtC::pKM3,* $tp^r$ | *mgtC* | (Maloney & Valvano, 2006) |
| ***Burkholderia thailandensis*** | | | |
| E264 | wild-type | wild-type | (Daligault et al., 2014) |
| MP1936 | *ΔmgtC::*Tn*10,* $tet^R$ | *mgtC* | This study |
| ***Klebsiella pneumoniae*** | | | |
| MKP103 | wild-type | wild-type | (Ramage et al., 2017) |
| KP5811 | *mgtC::*T30*,* $cm^R$ | *mgtC* | (Ramage et al., 2017) |

| | | | |
|---|---|---|---|
| ***Photorhabdus luminescens*** | | | |
| TT01 | wild-type | wild-type | (Duchaud et al., 2003) |
| RB101 | *ΔmgtC::km*$^{R}$ | *mgtC* | This study |
| ***Pseudomonas aeruginosa*** | | | |
| PAO1 | wild-type | wild-type | (Stover et al., 2000) |
| PW8814 | *mgtC::phoA-tet*$^{R}$ | *mgtC* | (Jacobs et al., 2003) |
| ***Salmonella bongori*** | | | |
| S3041 | wild-type | wild-type | (Boyd et al., 1993) |
| RB52 | *ΔmgtC::km*$^{R}$ | *mgtC* | This study |
| ***Salmonella enterica* serovar Typhimurium** | | | |
| 14028s | wild-type | wild-type | (Fields et al., 1986) |
| EL4 | *ΔmgtC* | *mgtC* | (Lee et al., 2013) |
| SL233 | $P_{rpsM}$-*mCherry*, at attTn*7* | N/A | This study |
| SL234 | *ΔmgtC* ($P_{rpsM}$-*mCherry*), at attTn*7* | N/A | This study |
| RB391 | *ΔmgtC* ($P_{mgtC\text{-}leader}$-*mgtC*-FLAG), at att*7* | N/A | This study |
| ***Sodalis praecaptivus*** | | | |
| ATCC BAA-2554 | wild-type | wild-type | (Chari et al., 2015) |
| RB61 | *ΔmgtC::km*$^{R}$ | *mgtC* | This study |
| ***Yersinia enterocolitica*** | | | |
| JB580v | wild-type | wild-type | (Walker et al., 2013) |
| MP2500 | *ΔmgtC::km*$^{R}$ | *mgtC* | This study |
| ***Yersinia pestis*** | | | |
| KIM6 | *pgm*$^{-}$ *pst*$^{+}$ *lcr*$^{-}$ *fra*$^{+}$ (pmT1, pCP1) | wild-type | (Staggs & Perry, 1991) |
| RB51 | KIM6 *ΔmgtC::km*$^{R}$ | *mgtC* | This study |
| **Plasmids** | | | |
| pKD4 | $rep_{R6K\gamma}$ *amp*$^{R}$ FRT-*km*$^{R}$-FRT | pKD4 | (Datsenko & Wanner, 2000) |
| pKD4-*tet*$^{R}$ | $rep_{R6K\gamma}$ *amp*$^{R}$ FRT-Tn*10*-FRT | pKD4-Tn*10* | (Pontes & Groisman, 2019) |
| pKD4-*apr*$^{R}$ | $rep_{R6K\gamma}$ *amp*$^{R}$ FRT- *apr*$^{R}$ -FRT | pKD4-*apr*$^{R}$ | (Pontes & Groisman, 2019) |
| pSIM5 | $rep_{pSC101}^{ts}$ *cm*$^{R}$ $P_{CI857}$-γβexo | pSIM5 | (Datta et al., 2006) |
| pSIM6 | $rep_{pSC101}^{ts}$ *amp*$^{R}$ $P_{CI857}$-γβexo | pSIM6 | (Datta et al., 2006) |
| pSIM19 | $rep_{pSC101}^{ts}$ spec$^{R}$ $P_{CI857}$-γβexo | pSIM19 | (Datta et al., 2006) |
| pAT02 | $rep_{RSF1010}$ amp$^{R}$ *lacI* P*tac*-RecAB | | (Tucker et al., 2014) |
| pAT03 | $rep_{RSF1010}$ amp$^{R}$ *lacI* P*tac*-FLP | | (Tucker et al., 2014) |
| pAOJ15 | $rep_{RK2}$ *amp*$^{R}$ *tetRA::tse2* | N/A | (Armbruster et al., 2017) |
| pAOJ15-*mgtC::km*$^{R}$ | $rep_{RK2}$ *amp*$^{R}$ *mgtC::km*$^{R}$ (*km*$^{R}$ cassette flanked by DNA sequence surrounding the *mgtC* gene from *Photorhabdus luminescens*) | pAOJ15_ *mgtC*::Km | This study |
| pGRG36-$P_{mgtC\text{-}leader}$-*mgtC*-FLAG | $rep_{pSC101}^{ts}$ *amp*$^{R}$ Tn*7*::( P*mgtC-leader-mgtC*-FLAG) | N/A | This study |
| pMRE-Tn7-166-P1 | $rep_{pSC101}^{ts}$ *amp*$^{R}$ Tn*7::*(*cm*$^{R}$ *tet*$^{R}$ *mCardinal*) | N/A | (Keller et al., 2021) |
| pTn7-mCherry-P1 | $rep_{pSC101}^{ts}$ *amp*$^{R}$ Tn*7*::( $P_{rpsM}$-*mCherry*) | N/A | This study |

| | | | |
|---|---|---|---|
| pAnl | rep$_{pMB1}$ *km*$^{R}$ *NLL-metRS* | pAnl | This study |
| pGFP | rep$_{p15A}$ *cm*$^{R}$ promoterless *gfp* vector control | pACYC-GFPc, pVector | (Pontes & Groisman, 2018) |
| pPstS-GFP | rep$_{p15A}$ *cm*$^{R}$ P*pstS*-*gfp* | pACYC-P*pstS*-GFPc | (Pontes & Groisman, 2018) |
| pBAD18 | rep$_{pMB1}$ *araC amp*$^{R}$ P*bad* | pBAD18, pBad | (Guzman et al., 1995) |
| pBAD18-GFP | rep$_{pMB1}$ *araC amp*$^{R}$ P*bad*- *gfp* | pBad-GFP | Lab plasmid |
| pBAD18-MgtC | rep$_{pMB1}$ *araC amp*$^{R}$ P*bad*- *mgtC* (*mgtC* from *Salmonella enterica*) | pBad-MgtC, pBad-MgtC$_{Se}$ | Lab plasmid |
| pBAD18-MgtC$_{Bt}$ | rep$_{pMB1}$ *araC amp*$^{R}$ P*bad*- *mgtC*$_{Bt}$ (*mgtC* from *Burholderia thailandensis*) | pBad-MgtC$_{Bt}$ | This study |
| p18 | rep$_{pMB1}$ *amp*$^{R}$ | p18, pVector | This study |
| p18-MgtC$_{Se}$ | rep$_{pMB1}$ *amp*$^{R}$ P*mgtC*$_{Se}$-*mgtC* (*mgtC* from *Salmonella enterica*) | p18-P*mgtC*$_{Se}$-MgtC, pMgtC$_{Se}$ | This study |
| p18-MgtC$_{Kp}$ | rep$_{pMB1}$ *amp*$^{R}$ P*mgtC*$_{Se}$-*mgtC*$_{Kp}$ (*mgtC* from *Klebsiella pneumoniae*) | p18-P*mgtC*$_{Se}$-MgtC$_{Kp}$, pMgtC$_{KP}$ | This study |
| pUHE-21–2-*lacI*$^{q}$ | rep$_{pMB1}$ *lacI*$^{q}$ *amp*$^{R}$; vector control | pUHE-21, pVector | (Soncini et al., 1995) |
| pUHE-*mgtC*$_{Bt}$*::*Tn*10* | rep$_{pMB1}$ *lacI*$^{q}$ *amp*$^{R}$ Tn*10* (*tet*$^{R}$ cassette flanked by DNA sequence surrounding the *mgtC* gene from *Burholderia thailandensis*) | N/A | This study |
| pUHE-*mgtC*$_{Ab}$*::apr*$^{R}$ | rep$_{pMB1}$ *lacI*$^{q}$ *amp*$^{R}$ Tn*10* (*apr*$^{R}$ cassette flanked by DNA sequence surrounding the *mgtC* gene from *Acinetobacter baumannii*) | N/A | This study |
| pUHE-MgtC | rep$_{pMB1}$ *lacI*$^{q}$ *amp*$^{R}$ P*lac*- *mgtC* (*mgtC* from *Salmonella enterica*) | pMgtC | (Chamnongpol & Groisman, 2002) |
| pUHE-MgtC$_{Phot}$ | rep$_{pMB1}$ *lacI*$^{q}$ *amp*$^{R}$ P*lac*- *mgtC*$_{Phot}$ (*mgtC* from *Photorhabdus luminescens*) | pMgtC$_{Phot}$ | This work |
| pUHE-MgtC$_{Soda}$ | rep$_{pMB1}$ *lacI*$^{q}$ *amp*$^{R}$ P*lac*- *mgtC*$_{Soda}$ (*mgtC* from *Sodalis praecaptivus*) | pMgtC$_{Soda}$ | This study |

*amp*$^{R}$ – ampicillin resistance; *apr*$^{R}$ – apramycin resistance; chloramphenicol resistance; *km*$^{R}$ – kanamycin resistance; *spec*$^{R}$ – spectinomycin resistance; *tet*$^{R}$ – tetracycline resistance; *tp*$^{R}$ – trimethoprim resistance.

### Table S2. Oligonucleotides sequences used in this study

| Name | Sequence (5'→ 3') | Purpose |
|---|---|---|
| 1547 | AAATTAACTATGAGAGGGACCACGGCGACGGCCAGC | Amplification of DNA regions flanking *Bukholderia thailandensi*s' *mgtC* homolog |
| 1548 | GAAGCAGCTCCAGCCTACACCCTGCCATGCCGAATGCGC | Amplification of DNA regions flanking *Bukholderia thailandensi*s' *mgtC* homolog |
| 1549 | TAAGGAGGATATTCATATGGTGACGCGCCGCGAGCCG | Amplification of DNA regions flanking *Bukholderia thailandensi*s' *mgtC* homolog |

| | | |
|---|---|---|
| 1550 | CAAGCTCAGCTAATTAAGCTCTAGCAAGTA TGACAGCGTG | Amplification of DNA regions flanking *Bukholderia thailandensi*s' *mgtC* homolog |
| 1551 | GTGTAGGCTGGAGCTGCTTC | Amplification of Tn*10* to generate *Bukholderia thailandensi*s' *mgtC::*Tn*10* allele |
| 1552 | CATATGAATATCCTCCTTA | Amplification of Tn*10* to generate *Bukholderia thailandensi*s' *mgtC::*Tn*10* allele |
| 145 | CAATTGTGAGCGGATAACAATTTC | Verification of *ΔmgtC::* Tn*10* allele from *Bukholderia thailandensi*s in pUHE-21 plasmid. |
| 146 | AATCCAGATGGAGTTCTGAGG | Verification of *ΔmgtC::* Tn*10* allele from *Bukholderia thailandensi*s in pUHE-21 plasmid. |
| 1553 | GACCACGGCGACGGCCAGC | Amplification of *mgtC*::Tn*10* to inactivate *Bukholderia thailandensi*s' *mgtC* gene |
| 1554 | CTAGCAAGTATGACAGCGTG | Amplification of *mgtC*::Tn*10* to inactivate *Bukholderia thailandensi*s' *mgtC* gene |
| 1716 | GTGCTGCTCAACGAGCACC | Verification of *ΔmgtC::*Tn*10* insertion in the *Bukholderia thailandensi*s' chromosome |
| 1717 | GAATGTATCAGTCCGCGCGC | Verification of *ΔmgtC::*Tn*10* insertion in the *Bukholderia thailandensi*s' chromosome |
| 143 | TGTCCAGATAGCCCAGTAGC | Verification of insertions generated with pKD4 |
| 493 | TCGAATTATTTAGGGTATAAATAGCACATG ATTATTCTACCAATGATTATTTCGTGTAGGC TGGAGCTGCTTC | Inactivation of *Yersinia pestis' mgtC (ΔmgtC::km*$^R$ generation) |
| 494 | TTGTCGTTTTATCTATTTATCGTTTTATCTAT TTATTGCTTAATTAATGACTCGCATATGAAT ATCCTCCTTA | Inactivation of *Yersinia pestis' mgtC (ΔmgtC::km*$^R$ generation) |
| 492 | TAGGTATATACCGTCATAT | Verification of *ΔmgtC::km*$^R$ insertion in the *Yersinia pestis'* chromosome |
| 2380 | ACATGATTATTCTACCAATGATTAATTCATG GAGAGAATCGTGTAGGCTGGAGCTGCTTC | Inactivation of *Yersinia enterocolitica's mgtC (ΔmgtC::km*$^R$ generation) |
| 2381 | TATTGACTGCTAATTTGAAAGATATATTAA TTCCCTAACCCATATGAATATCCTCCTTAG | Inactivation of *Yersinia enterocolitica's mgtC (ΔmgtC::km*$^R$ generation) |
| 2383 | TCAGCACAATGACCTGTGT | Verification of *ΔmgtC::km*$^R$ insertion in the *Yersinia enterocolitica's* chromosome |
| 490 | TCGTGTGCTAAATATAGCACGTACTTATTCT TCCAGAAAAACATATGAATATCCTCCTTA | Inactivation of *Salmonella bongori's mgtC (ΔmgtC::km*$^R$ generation) |
| 491 | TATACGCCTGGCGTAATGTTGAAATTGAAT AAAAAAACCAGTGTAGGCTGGAGCTGCTTC | Inactivation of *Salmonella bongori's mgtC (ΔmgtC::km*$^R$ generation) |
| 489 | CTCCACCGTCAACACGACGC | Verification of *ΔmgtC::km*$^R$ insertion in the *Salmonella bongori's* chromosome |

| | | |
|---|---|---|
| 496 | CGTGCGCAAAAAGAAAACGTAATTACGCCAACCATGAGGAAAGCGTTCATATGAATATCCTCCTTA | Inactivation of *Sodalis praecaptivus' mgtC* ($\Delta mgtC::km^R$ generation) |
| 497 | CTTTCCCGCCTGTCGCCGGATGGGGGTTCATTCATAGTCATCTCCTGCGTGTAGGCTGGAGCTGCTTC | Inactivation of *Sodalis praecaptivus' mgtC* ($\Delta mgtC::km^R$ generation) |
| 495 | ATTACACGACGCGAATAAC | Verification of $\Delta mgtC::km^R$ insertion in the *Sodalis praecaptivus'* chromosome |
| 567 | TATTCTCCCGGCATCTCTC | Verification of $\Delta mgtC::km^R$ insertion in the *Sodalis praecaptivus'* chromosome |
| 1 | GTGTAGGCTGGAGCTGCTTC | Amplification of $km^R$ gene from pKD4 for construction of *Photorhabdus luminescens'* $\Delta mgtC::km^R$ allele |
| 2 | CATATGAATATCCTCCTTA | Amplification of $km^R$ gene from pKD4 for construction of *Photorhabdus luminescens'* $\Delta mgtC::km^R$ allele |
| 528 | TGAATTCGAGCTCGGTACCCGGGCTTTTCTATCATGAGTTAATGCAG | Amplification of DNA regions flanking *Photorhabdus luminescens' mgtC* homolog |
| 529 | TTCGAAGCAGCTCCAGCCTACACCATGACATTCTCCATACAACC | Amplification of DNA regions flanking *Photorhabdus luminescens' mgtC* homolog |
| 530 | GAACTAAGGAGGATATTCATATGTCTGAACTACCGATCTGAAAAATCT | Amplification of DNA regions flanking *Photorhabdus luminescens' mgtC* homolog |
| 531 | TATGACCATGATTACGCCAAGCTCGACATTCTCCATCAAGCAG | Amplification of DNA regions flanking *Photorhabdus luminescens' mgtC* homolog |
| 144 | CTGGATGATCCTCCAGCGCG | Verification of pAOJ15-$mgtC::km^R$ insertion in the *Photorhabdus luminiscens'* chromosome |
| 568 | CACCTATGCATAAAAAGGTTGC | Verification of $\Delta mgtC::km^R$ insertion in the *Photorhabdus luminiscens'* chromosome |
| 569 | GTAATATCATCACGCTTTCATCACC | Verification of $\Delta mgtC::km^R$ insertion in the *Photorhabdus luminiscens'* chromosome |
| 277 | CACACTTTGCTATGCCAT | pBAD18 sequencing |
| 278 | CGCTACTGCCGCCAGGCA | pBAD18 sequencing |
| 2424 | TGAGCGCTTGTTTCGGCGT | Construction of p18 plasmids |
| 2425 | ATGGAGGAACGTATGTTAATGTT | Construction of p18-MgtC$_{Sent}$ |
| 2426 | ATGGTATTTCCTTATATGACTCATTTACT | Construction of p18-MgtC$_{Kpne}$ |
| 2428 | CCACGCCGAAACAAGCGCTCAGAGAGAAGATTTTCAGCCTG | Construction of p18 plasmids |
| 2429 | CCACGCCGAAACAAGCGCTCAGGAGAAGGTCACCAGTATCA | Amplification of MgtC$_{Sent}$ cis-regulatory elements |
| 2430 | GGAAACATTAACATACGTTCCTCCATTTTTTCTGGAAGAATAAGTACGT | Amplification of MgtC$_{Sent}$ cis-regulatory elements |
| 2431 | AGTAAATGAGTCATATAAGGAAATACCATTTTTTCTGGAAGAATAAGTACGT | Amplification of MgtC$_{Sent}$ cis-regulatory elements |

| | | |
|---|---|---|
| 2322 | GTTTTTTTGGGCTAGCGAATTCGCATTCGGCATGGCAGGAC | Amplification of *mgtC* from *B. thailandensis* for cloning into pBAD18 |
| 2323 | CTCATCCGCCAAAACAGCCAAGCTTGTCGTGAAGGCGATCGCGAT | Amplification of *mgtC* from *B. thailandensis* for cloning into pBAD18 |
| 146 | AATCCAGATGGAGTTCTGAGG | pUHE-21–2-*lacI*$^{q}$ sequencing |
| 153 | GTCTCATGAGCGGATACATAT | pUHE-21–2-*lacI*$^{q}$ sequencing |
| 2259 | AGGAGAAATTAACTATGAGAGGATGAGGAAAGCGTTATGGTAT | Cloning of *mgtC* from *Sodalis praecaptivitus* into pUHE-21 |
| 2260 | AGCTAATTAAGCTTGGCTGCATGGGGGTTCATTCATAGTC | Cloning of *mgtC* from *Sodalis praecaptivitus* into pUHE-21 |
| 2261 | AGGAGAAATTAACTATGAGAGGATTGGTTGTATGGAGAATGTCA | Cloning of *mgtC* from *Photorhabdus luminiscens* into pUHE-21 |
| 2262 | AGCTAATTAAGCTTGGCTGCTCAGATCGGTAGTTCAGAAG | Cloning of *mgtC* from *Photorhabdus luminiscens* into pUHE-21 |
| 1291 | TCAGATCCCGGGTCAATAGCCTTTGCTCCATGATGTACTG | Construction of pGRG36-$P_{mgtC\text{-}leader}$-*mgtC*-FLAG |
| 1301 | CTTGTCGTCATCGTCCTTGTAGTCCATTTTTTCTGGAAGAATAAGTACGTG | Construction of pGRG36-$P_{mgtC\text{-}leader}$-*mgtC*-FLAG |
| 1302 | ACAAGGACGATGACGACAAGGACAAGGACAACACCGTGC | Construction of pGRG36-$P_{mgtC\text{-}leader}$-*mgtC*-FLAG |
| 1294 | GTGGCGCGCCTCCTAGGTGCTTACTTCTCTGCGCTTCTCAG | Construction of pGRG36-$P_{mgtC\text{-}leader}$-*mgtC*-FLAG |
| 785 | TTGTCTCATGAGCGGATACA | Construction of pMRE-Tn7-mCherry-P1 |
| 786 | ATGATCGACCGAGACAGGC | Construction of pMRE-Tn7-mCherry-P1 |
| 3028 | GCCTGTCTCGGTCGATCATCGTGGCTTACTAGGATCCGA | Construction of pMRE-Tn7-mCherry-P1 |
| 3029 | TGTATCCGCTCATGAGACAAATCATGGTCCTGCTTGCTT | Construction of pMRE-Tn7-mCherry-P1 |
| 92 | TAGCGTCGTAAGCTAATACGA | Sequencing inserts cloned into pGRG36/pMER-Tn*7* |
| 93 | CAGTCCAGTTATGCTGTGA | Sequencing inserts cloned into pGRG36/pMER-Tn*7* |
| 156 | GGCACCGACGTTGACCAGC | Verification of *mgtC* insertion at the Tn*7* attachment site |
| 157 | CGTTCAGGCTGGCTACAGC | Verification of *mgtC* insertion at the Tn*7* attachment site |
| 1296 | TGCTTAATACCGGCGATAGC | Verification of *mgtC* insertion at the Tn*7* attachment site |
| 1299 | GCGACAAGGAAATCTTCG | Verification of *mgtC* insertion at the Tn*7* attachment site |
| 2336 | CCGGGAGCTGCATGTGTCAGAGG | Verify pAnl by Sanger DNA sequencing |
| 2337 | ACGTGCAGTCGATGATAAG | Verify pAnl by Sanger DNA sequencing |
| 2263 | GCAACGGGATTGAGGGATAG | qPCR for *Sodalis praecaptivus' mgtC* |
| 2264 | TGCGGACATCATGGCTTATC | qPCR for *Sodalis praecaptivus' mgtC* |
| 2265 | GGTAAATCCAACCCGGTTACT | qPCR for *Sodalis praecaptivus' ompA* |

| | | |
|---|---|---|
| 2266 | TGCCTTTCACTTCGATCTCTAC | qPCR for *Sodalis praecaptivus' ompA* |
| 2269 | GTCAACGTATGGCAGGTTTAAG | qPCR *Photorhabdus luminiscens' mgtC* |
| 2270 | CACTATCGGGTGAAGTTGTTATTG | qPCR *Photorhabdus luminiscens' mgtC* |
| 2271 | CAATCGGTAATGGCCCTACTC | qPCR *Photorhabdus luminiscens' ompA* |
| 2272 | GCCTAACCAGTCGTAACCTATTT | qPCR *Photorhabdus luminiscens' ompA* |
| 2553 | TGTGTAGGCTGGAGCTGCTT | Amplification of $apr^R$ to generate *Acinetobacter baumannii*' *mgtC::apr*$^R$ allele |
| 2554 | CCTCCTTAGTTCCTATTCCGA | Amplification of $apr^R$ to generate *Acinetobacter baumannii*' *mgtC::apr*$^R$ allele |
| 2555 | AATTAACTATGAGAGGATCCGTGGATGAAG AGTTAATTGCA | Amplification of DNA regions flanking *Acinetobacter baumannii*' *mgtC* homolog |
| 2556 | AAGCAGCTCCAGCCTACACACACCTATTGT TTTATTGCCA | Amplification of DNA regions flanking *Acinetobacter baumannii*' *mgtC* homolog |
| 2557 | TCGGAATAGGAACTAAGGAGGAAATGATG AAGCGCAAACTC | Amplification of DNA regions flanking *Acinetobacter baumannii*' *mgtC* homolog |
| 2558 | TCCAAGCTCAGCTAATTAAGCTTGCAACCC TCACAAGCTT | Amplification of DNA regions flanking *Acinetobacter baumannii*' *mgtC* homolog |
| 2559 | AGCTTAATTAGCTGAGCTTGGA | Amplification of *mgtC::apr*$^R$ to inactivate *Acinetobacter baumannii*' *mgtC* gene |
| 2561 | GGGTCCTTTAGCAAGCT | Amplification of *ΔmgtC::apr*$^R$ from pUHE-*mgtC*$_{Ab}$*::apr*$^R$ |
| 2562 | AGCGGCATTATTTAC | Amplification of *ΔmgtC::apr*$^R$ from pUHE-*mgtC*$_{Ab}$*::apr*$^R$ |
| 2563 | GTCCATATGATCCGTACA | Verification of *ΔmgtC::apr*$^R$ insertion in the *Acinetobacter baumannii*' chromosome |
| 2564 | CAAGTTCATCCTGATCCA | Verification of *ΔmgtC::apr*$^R$ insertion in the *Acinetobacter baumannii*' chromosome |

### Supplemental Materials and Methods

#### Recombineering protocols

Phusion High-Fidelity DNA Polymerase (New England Biolabs) was used in polymerase chain reactions (PCRs) to construct *ΔmgtC::km*$^R$ alleles. Specifically, plasmid pKD4 (Datsenko & Wanner, 2000) was used as DNA template in PCR reactions with 490/491, 493/494, and 496/497 to generate *ΔmgtC::km*$^R$ alleles for inactivation of *S. bongori*, *S. praecaptivus* , *Y. enterocolitica* and *Y. pestis mgtC* homologs, respectively. PCR reactions were separated using agarose gel electrophoresis and bands corresponding to the PCR products were purified using the Monarch DNA Gel Extraction Kit (New England Biolabs).

*Salmonella bongori* harboring plasmid pSIM6 (Datta et al., 2006) was grown overnight in LB broth supplemented with 100 µg/ mL of ampicillin at 30°C and 250 rpm shaking water bath. The culture was diluted (1:100) in 30 mL of the same medium and grown under the same conditions for approximately 2.5 h ($OD_{600}$ ~0.35-0.4). The recombineering functions encoded in pSIM6 were induced by transferring the culture flask to a 42°C and 250 rpm shaking water bath for 20 min (final $OD_{600}$ ~0.6-0.75). The culture was then transferred to a 50 v conical tube, collected by centrifugation (4°C, 7,000 rpms for 2.5 min), washed three times with ice-cold type-1 water, and resuspended in 150 µl of ice-cold type-1 water. Insertional inactivation of *mgtC* was achieved by electroporating 70 µl of these cells with purified, PCR-generated *ΔmgtC::km*$^R$ allele, allowing the cells to recover for 2 h at 37°C and 250 rpm and selecting for recombinants on LB agar plates supplemented with 50 µg/ mL of kanamycin. The location of insert in recombinant bacteria was verified using OneTaq Polymerase (New England Biolabs) and primer pair 143/489.

*Sodalis praecaptivus* harboring plasmid pSIM6 (Datta et al., 2006) was grown overnight in LB broth supplemented with 10 mM $MgCl_2$ and 100 µg/mL of ampicillin at 30°C and shaking at 250 rpm. Cells were diluted (1:20) in 60 mL of the same medium and grown for approximately 9 h ($OD_{600}$ ~0.5-0.55). To express recombineering functions encoded in pSIM6, the flask was transferred to a 42°C and 250 rpm shaking water bath for 1 h (final $OD_{600}$ ~0.6). Following induction of recombineering functions, cultures were transferred to two 50 mL conical tubes, collected by centrifugation (4°C, 7,000 rpms for 2.5 min), washed three times with ice-cold type-1 water, and resuspended in a total combined volume of 150 µl of ice-cold type-1 water. Insertional inactivation of *mgtC* was achieved by electroporating 70 µl of these cells with purified, PCR-generated *ΔmgtC::km*$^R$ allele, allowing cells to recover for 2 h at 30°C and 250 rpm and selecting for recombinants on LB agar plates supplemented with 50 µg/mL of kanamycin. The location of insert in recombinant bacteria was verified using OneTaq Polymerase (New England Biolabs) and primer pair 143/495.

*Yersinia enterocolitica* harboring plasmid pSIM19 (Datta et al., 2006) was grown overnight in LB broth supplemented with 100 µg/mL of spectinomycin at 30°C and shaking at 250 rpm. Cells were diluted (1:60) in 60 mL of the same medium and grown for approximately 4 h (OD600 ~0.35-0.4). The induction of recombineering genes in the pSIM19 plasmid was achieved by transferring the culture flask to a 42°C and 250 rpm shaking water bath for 45 min (final $OD_{600}$ ~0.6-0.75). Recombinogenic cells were collected by centrifugation (4°C, 7,000 rpms for 2.5 min), washed three times with ice-cold type-1 water, and resuspended in 150 µl of ice-cold type-1 water. Insertional inactivation of *mgtC* was accomplished by electroporating 70 µl of these cells with purified PCR-generated *ΔmgtC::km*$^R$ allele, allowing cells to recover for 2 h at 30°C and 250 rpm and selecting for recombinants on LB agar plates supplemented with 50 µg/mL of kanamycin. The location of insert in recombinant bacteria was verified using OneTaq Polymerase (New England Biolabs) and primer pair 143/2383.

*Yersinia pestis* harboring plasmid pSIM5 (Datta et al., 2006) was grown overnight in BHI broth supplemented with 20 µg/mL of chloramphenicol at 30°C and shaking at 250 rpm. Cells were diluted (1:20) in 60 mL of the same medium and grown for approximately 7 h (OD600 ~0.35-0.4). The induction of recombineering genes in the pSIM5 plasmid was achieved by transferring the culture flask to a 42°C and 250 rpm shaking water bath for 45 min (final $OD_{600}$ ~0.6-0.75). Recombinogenic cells were collected by centrifugation (4°C, 7,000 rpms for 2.5 min), washed three times with ice-cold type-1 water, and resuspended in 150 µl of ice-cold type-1 water. Insertional inactivation of *mgtC* was accomplished by electroporating 70 µl of these cells with purified PCR-generated *ΔmgtC::km*$^R$ allele, allowing cells to recover for 2 h at 30°C and 250 rpm and selecting for recombinants on BHI agar plates supplemented with 50 µg/mL of kanamycin. The location of insert in recombinant bacteria was verified using OneTaq Polymerase (New England Biolabs) and primer pair 143/492.

**Conjugation between *Escherichia coli* and *Photorhabdus luminescens* and isolation of *Photorhabdus luminescens ΔmgtC::km$^R$* mutant**

Phusion High-Fidelity DNA Polymerase (New England Biolabs) was used in PCRs with primer pairs 1/2 and plasmid pKD4 as template(Datsenko & Wanner, 2000), and primer pairs 528/529 and 530/531 and *P. luminescens* genomic DNA as template. PCR reactions were separated using agarose gel electrophoresis and bands corresponding to the PCR products were purified using the Monarch DNA Gel Extraction Kit (New England Biolabs). Purified PCR products were ligated into plasmid pAOJ15(Armbruster et al., 2017), previously digested with restriction enzymes BamHI and HindIII (New England BioLabs), using NEBuilder HiFi DNA Assembly Cloning Kit (New England BioLabs). The assembly was transformed into λpir$^+$ *E. coli* cells for plasmid propagation and sequencing.

Constructs with the correct sequence (designated pAOJ15-*mgtC::km$^R$*) were electroporated into *E. coli* BW29427, which has a mutation in the *dapA* gene. *Escherichia coli* BW29427 harboring pAOJ15-*mgtC::km$^R$* (donor strain) was cultured overnight at 250 rpms and 37 ˚C in LB medium supplemented with 50 µg/mL kanamycin and 60 µg/mL diaminopimelic acid (DAP). Wild-type *P. luminiscens* TT01 (recipient strain) was cultured overnight in BHI medium at 30˚C. A sample (200 µl) of each culture was sub-cultured into 20 mL of pre-warmed LB and incubated at either 37˚C (donor strain) or 30°C (recipient strain) at 250 rpms for 2 h ($OD_{600}$ ~0.3-0.4). Cultures were then centrifuged (3,000 rpm for 5 min), cells were washed once with BHI, and resuspended in PBS. The donor and recipient strains were then combined at a 1:5 (donor:recipient) ratio, centrifuged (3,000 rpm for 5 min), and gently resuspended in 50 µl of PBS. Cells mixture was spotted onto BHI-agar plates supplemented with 0.3 mM DAP and incubated at 30°C overnight. The conjugation mixture was scraped from the plates with a sterile pipet tip and transferred with a 1.5 mL microcentrifuge tube. The tube was centrifuged (3,000 rpm for 5 min), washed twice with PBS, and spread onto BHI agar plates containing 35 µg/mL kanamycin. Resulting colonies were re-plated onto LB agar containing 100 µg/mL ampicillin and 50 µg/ mL kanamycin. OneTaq DNA Polymerase (New England Biolabs) was used with primer pair 144/569 to verify pAOJ15-*mgtC::km$^R$* insertion and merodiploidy of *P. luminiscens* colonies. Merodiploids were cultured overnight in BHI broth with 35 µg/mL kanamycin at 30˚C and shaking at 250 rpm. Cells were diluted (1:100) in 2 mL of the same medium and grown for approximately 3 h ($OD_{600}$ ~0.6) at 30°C. 1 µg/mL anhydrotetracycline (Sigma Aldrich) was added to the cultures to promote the expression of the Tse2 toxin and counter select cells harboring pAOJ15-*mgtC::km$^R$* replicating with the chromosome or as an episome. Resulting colonies were re-streak onto LB agar containing anhydrotetracycline and 35 µg/mL kanamycin and incubated overnight at 30˚C. Verification for the exchange of *mgtC* coding sequence with the kanamycin resistance cassette was confirmed by PCR using OneTaq Polymerase (New England Biolabs) and primer pair 568/569.

**Construction of *ΔmgtC::*Tn*10* mutant strain of *Burholderia thailandensis***

Phusion High-Fidelity DNA Polymerase (New England Biolabs) was used in PCRs with primer pairs 1547/1548 and 1549/1550 and *Burkholderia thailandensis* genomic DNA as template, and 1551/1552 and plasmid pKD4-*tet$^R$* as template(Pontes & Groisman, 2019). PCR reactions were separated using agarose gel electrophoresis and bands corresponding to the PCR products were purified using the Monarch DNA Gel Extraction Kit (New England Biolabs). Purified PCR products were ligated into pUHE-21 plasmid (Soncini et al. 1995), previously digested with BamHI and HindIII, using NEBuilder HiFi DNA Assembly (New England Biolabs). Assembly was transformed into *E. coli* cells, which were selected on LB plates containing ampicillin and tetracycline. The *ΔmgtC::Tn10* allele was verified by DNA sequencing with primers 145 and 146. Plasmids with correct construct were designated pUHE-

*mgtC*$_{Bt}$*::Tn10* and saved. Phusion High-Fidelity DNA Polymerase (New England Biolabs) was used in a PCR with primer pair 1553/1554 and pUHE-*mgtC*$_{Bt}$*::Tn10* as template. PCR was separated using agarose gel electrophoresis and the band corresponding to the *ΔmgtC::Tn10* product was purified using the Monarch DNA Gel Extraction Kit (New England Biolabs). Purified PCR product was incubated with *B. thailandensis* cells that were rendered naturally competent and recombined with the purified PCR product as described(Thongdee et al., 2008). Recombinant bacteria were selected on LSLB plates containing tetracycline. Chromosomal insertion in tetracycline resistant colonies were verified using PCR with primers 1716/1717 and Sanger DNA sequencing.

**Construction of *ΔmgtC* mutant strain of *Acinetobacter baumannii***

The *mgtC* deletion mutant was generated using a recombineering approach (Tucker et al., 2014). Briefly, RepliQa HiFi DNA Polymerase (Quantabio) was used in PCRs with primer pairs 2553/2554 and pKD4-*apr*$^{R}$ (Pontes & Groisman, 2019) as template, and primer pairs 2555/2556 or 2557/2558 and *A. baumannii* genomic DNA as template. PCR reactions were separated using agarose gel electrophoresis and bands corresponding to the PCR products were purified using the Monarch DNA Gel Extraction Kit (New England Biolabs). Purified PCR products were ligated into pUHE-21 plasmid (Soncini et al. 1995), previously digested with BamHI and HindIII, using NEBuilder HiFi DNA Assembly (New England Biolabs). The *ΔmgtC::apr*$^{R}$ allele was verified by whole plasmid sequencing. Plasmids with correct construct were designated pUHE-*mgtC*$_{Ab}$*::apr*$^{R}$ and saved. RepliQa HiFi DNA Polymerase (Quantabio) was used in PCR with primer pairs 2561/2562 and pUHE-*mgtC*$_{Ab}$*::apr*$^{R}$ as template. PCR was separated using agarose gel electrophoresis and the band corresponding to the *ΔmgtC::apr*$^{R}$ product was purified using the Monarch DNA Gel Extraction Kit (New England Biolabs). Approximately 5 µg of the purified PCR product was electroporated into *A. baumannii* cells carrying the pAT02 plasmid, which expresses the RecAb recombination system under IPTG induction (2 mM). Transformants were selected on LB agar containing apramycin. Transformants were screened by colony PCR using primer pairs 2563/2564, which flank the *ΔmgtC::apr*$^{R}$ insertion point. After curing pAT02 the *ΔmgtC::apr*$^{R}$ strains were transformed with pAT03, which expresses the FLP recombinase from an IPTG induction. Removal of the apramycin resistance gene and confirmation of sensitivity by PCR using primers 2553/2554 to detect its presence.

**Construction of *Salmonella enterica* strains harboring insertions at the Tn*7* attachment site**

*Salmonella enterica* strains were electroporated with plasmids pTn7-mCherry-P1 or pGRG36-P*mgtC-leader-mgtC*-FLAG. Transformants were recovered at 30°C on LB plates containing ampicillin and 10 mM glucose. Single colonies were grown for 6 h at 30°C in MOPS broth containing 25 mM arabinose as the sole carbon source. 20 µl of culture were spotted and streaked on LB plates containing 0.1% arabinose. Single colonies were recovered following overnight incubation at 42°C. Single colonies were then screened for the loss of plasmid (ampicillin sensitivity) and the presence of transposed DNA at the Tn*7* attachment site. Specifically, the expression of mCherry was verified by fluorescence (excitation 587 nm/emission 610 nm) using a SpectraMax i3x (Molecular Devices) plate reader. Ampicillin-sensitive clones that harbored pGRG36-P*mgtC-leader-mgtC*-FLAG were screened by colony PCR using OneTaq (New England Biolabs) and primer pairs 156/1299 and 157/1296.

**Construction of plasmids pUHE-MgtC$_{Phot}$ and pUHE-MgtC$_{Soda}$**

RepliQa HiFi DNA Polymerase (Quantabio) was used in PCRs with primer pairs 2259/2260 and *S. praecaptivus* genomic DNA as template; and primer pairs 2261/2262 and *P. luminescens* genomic DNA as template. PCR reactions were separated using agarose gel electrophoresis and bands corresponding to the PCR products were purified using the Monarch DNA Gel Extraction

Kit (New England Biolabs). Purified PCR products were ligated into pUHE-21 plasmid (Soncini et al., 1995), previously digested with BamHI and PstI, using NEBuilder HiFi DNA Assembly (New England Biolabs). Assembly was transformed into *E. coli* cells, which were selected on LB plates containing ampicillin. The integrity of the constructs was verified by whole plasmid sequencing.

**Construction of plasmids p18, p18-P*mgtC$_{Se}$-mgtC$_{Se}$*, p18-P*mgtC$_{Se}$-mgtC$_{Kp}$*, and pBAD18-*mgtC$_{Bt}$***

RepliQa HiFi DNA Polymerase (Quantabio) was used in PCR reactions with primer pairs 2424/2428 and plasmid as template pBAD18 (Guzman et al., 1995), primer pairs 2424/2425 and plasmid pBAD18-MgtC$_{Se}$ (lab plasmid) DNA as template, primer pairs 2424/2426 and plasmid pBAD18-MgtC$_{Kp}$ (lab plasmid) DNA as template, primer pairs 2585/2586 and p18-MgtC$_{Se}$ (p18-P*mgtC$_{Se}$-mgtC$_{Se}$*) DNA as template, 2322/2223 and *B. thailandensis* E264 genomic DNA as template, 2429/2430 or 2429/2431 and *S. enterica* 14028s genomic DNA as template. PCR reactions were digested with DpnI (New England Biolabs), separated using agarose gel electrophoresis, and bands corresponding to the PCR products were purified using the Monarch DNA Gel Extraction Kit (New England Biolabs). Plasmid p18 was generated by transforming the PCR product obtained with primers 2424/2428 into *E. coli*. Plasmid p18-MgtC$_{Se}$ (p18-P*mgtC$_{Se}$-mgtC$_{Se}$*) was generated by transforming *E. coli* with a NEBuilder HiFi DNA Assembly (New England Biolabs) of PCR fragments generated with primers 2424/2425 and 2429/2430. Plasmid p18-MgtC$_{Kp}$ (p18-P*mgtC$_{Se}$-mgtC$_{Kp}$*) was generated by transforming *E. coli* with a NEBuilder HiFi DNA Assembly (New England Biolabs) of PCR fragments generated with primers 2424/2426 and 2429/2431. Plasmid pBAD18-MgtC$_{Bt}$ was generated by transforming *E. coli* with a T4 DNA ligase (New England Biolabs) reaction containing PCR fragment generated with primers 2322/2323 and plasmid pBAD18, both of which had been previously digested with EcoRI and BamHI. The integrity of the constructs was verified by whole plasmid sequencing.

**Construction of plasmid pAnl**

Plasmid pAM1 (Mahdavi et al., 2014) was digested with restriction enzymes EheI and FspAI (Thermo Fisher Scientific). The digestion was separated using agarose gel electrophoresis and the band corresponding to the plasmid backbone was purified using the Monarch DNA Gel Extraction Kit (New England Biolabs). The purified plasmid was self-ligated using T4 ligase (New England Biolabs). The ligation point was verified by Sanger DNA sequencing using primers 2336 and 2337.

**Construction of plasmid pGRG36-P*mgtC-leader-mgtC*-FLAG and pMRE-Tn7-mCherry-P1**

Phusion High-Fidelity DNA Polymerase (New England Biolabs) was used in a PCR with primer pair 1291/1301 and pGFP303 (Lee and Groisman, 2012) as template, and primer pair 1302/1294 and pUHE-*mgtC*-FLAG-BID (lab plasmid) as template. PCRs were separated using agarose gel electrophoresis and bands corresponding to the PCR products were purified using the Monarch DNA Gel Extraction Kit (New England Biolabs). Purified PCR products were ligated into plasmid pGRG36 (McKenzie and Craig, 2006), previously digested with restriction enzymes NotI and XhoI (New England BioLabs), using NEBuilder HiFi DNA Assembly Cloning Kit (New England BioLabs) to produce plasmid pGRG36-P$_{mgtC\text{-}leader}$-*mgtC*-FLAG.

Phusion High-Fidelity DNA Polymerase (New England Biolabs) was used in a PCR with primer pair 785/786 and pFPV25.1-mCherry (a lab plasmid harboring a transcriptional fusion between the *S. enterica rpsM* promoter and a *mCherry* gene) as template. Phusion High-Fidelity DNA Polymerase (New England Biolabs) was used in a PCR with primer pair 3028/3029 and plasmid pMRE-Tn7-166-P1 (Keller et al., 2021). PCR reactions were separated using agarose gel electrophoresis, and bands corresponding to the PCR products were purified using the Monarch DNA Gel Extraction Kit (New England Biolabs). Purified PCR products were assembled using NEBuilder HiFi DNA Assembly Cloning Kit (New England BioLabs) to produce plasmid pMRE-Tn7-mCherry-P1. All assembly reactions were transformed into *E. coli* cells, which were recovered at 30°C on LB plates containing ampicillin and 10 mM glucose, for plasmid propagation and sequencing.

**Measuring bacterial growth**

Physiological measurements of bacterial growth were carried out in MOPS medium (Neidhardt et al., 1974) lacking $CaCl_2$ and supplemented with 22 mM glucose, 0.1% casein amino acids, and the indicated concentrations of $MgCl_2$ and $K_2HPO_4$. For *A. baumannii*, 22 mM glucose was replaced with 38 mM glycerol. For *P. luminescens*, *S. praecaptivus*, *Y. enterocolitica* and *Y. pestis*, this medium was further supplemented with the following amino acid mixture: 3.2 mM of alanine, glycine, leucine, glutamate and serine; 2.4 mM glutamine, isoleucine and valine; 1.6 mM arginine, asparagine, aspartate, lysine, phenylalanine, proline, threonine, and methionine; 0.8 mM histidine and tyrosine, and 0.4 mM cysteine and tryptophan. In these experiments, bacteria were propagated to the stationary phase in this MOPS medium containing 10 mM $MgCl_2$ and 2 mM $K_2HPO_4$. Stationary phase cells were washed three times with 1x MOPS salts lacking $MgCl_2$ and $K_2HPO_4$, diluted 1:100 into fresh MOPS medium containing 10 µM $MgCl_2$ and either 500 µM or no exogenous $K_2HPO_4$, and subsequently aliquoted into clear 96-well plates (Corning). Two drops of mineral oil were used to seal the wells and prevent evaporation. Cultures were propagated at 30°C (*B. cenocepacia, B. thailandensis*, *P. luminescens*, *S. praecaptivus*, *Y. enterocolitica* and *Y. pestis*) or 37°C (*A. baumannii*, *E. coli, K. pneumoniae*, *P. aeruginosa*, *S. bongori*, *S. enterica*) with auto-mixing in a SpectraMax i3x (Molecular Devices) plate reader. The optical density at 600 nm ($OD_{600}$) was measured for each well at regular time intervals. Growth was inferred as the increase in $OD_{600}$ over time.

**mRNA extraction and quantification**

*Photorhabdus luminescens* and *S. praecaptivus* were grown at 30°C and 250 rpms in MOPS medium containing 10 mM $MgCl_2$ and 2 mM $K_2HPO_4$. Stationary phase cultures were diluted 1:100 into fresh medium and propagated to an $OD_{600}$ of 0.4-0.5. Cells were collected by centrifugation (14,000 x g for 1 min), washed three times with 1x MOPS salts lacking $MgCl_2$ and $K_2HPO_4$, and resuspended in this MOPS salts solution at $1/100^{th}$ of the original culture volume. These cells were then used to inoculate fresh MOPS medium containing 2 mM $K_2HPO_4$ and either 10 mM or no $MgCl_2$, to yield cultures with an $OD_{600}$ of 0.4-0.5. Cells were propagated at 30°C and 250 rpms for 2 h. RNA was stabilized using RNAprotect Bacteria (Qiagen) and extracted using the RNeasy Kit (Qiagen). cDNA was synthesized from RNA samples using the SuperScript VILO cDNA synthesis kit (Thermo Fisher Scientific). Relative amounts of cDNA were determined by quantitative PCR (qPCR) using primer pairs specific to *ompA* and *mgtC* (Table S2), and a standard curve generated from serially diluted genomic DNA from *P. luminescens* or *S. praecaptivus* (Pontes et al., 2011). qPCR reactions were performed using Fast SYBR Green master mix (Thermo Fisher Scientific) in an QuantStudio 7 Flex Real-Time PCR detection system (Thermo Fisher Scientific).

**Nucleotide sequences used in phylogenetic analyses**

16S rDNA sequences: STM14_0293, FU841_02320, b0201, AL524_00665, K7R23_00020, QJR40_00080, CHQ57_00385, FOB40_10380, SSYIS1_03910, Sant_r0262, H8F46_03010, AM461_18970, A6V27_02945, GO998_01710, RSOE_15460, BG16_4297, BTH_II2047,

BCAM2492, AL472_10800, SF4435, DJY80_01120, PP_16SA, PA0668.1, BFV67_01110, AM380_01395, CG030_04105, AM402_16995, GH767_00015, CRN74_01230, SMDB11_rRNA_20, DSM2777_01660, CS875_16035, B3286c2_0624, CBF16_16925, PluTT01m_RS22565, *rrsS* gene within *Mycobacterium tuberculosis* H37Rv genome ( genome sequence accession number AL123456), MMARE11_RS19305, A1S_r01.

*mgtC/yhiD* sequences: STM14_4538, FU841_12190, b3508, AL524_05215, QJR40_08400, CHQ57_02780, FOB40_00290, SSYIS1_02730, Sant_3759, AM461_10480, A6V27_09745, GO998_09210, RSOE_15605, BG16_4150, BTH_II2176, BCAM0411, AL472_22835, DJY80_09005, PP_3244, PA4635, BFV67_09760, AM380_19320, CG030_17870, AM402_09585, GH767_04830, CRN74_09755, SMDB11_2322, DSM2777_06170, CS875_16645, B3286c2_0751, CBF16_09930, PluTT01m_RS09545, Rv1811, MMARE11_RS12855, J5F57_002221, NX108_RS23335, LUM72_003877, RHV78_001718, WHU19_RS22485, MML20_003046, RHV70_002104, EPS86_23855, WHU19_22485, EPS86_RS23850, 1S_2069.

### Supplemental Material References


Armbruster, C. E., Forsyth-DeOrnellas, V., Johnson, A. O., Smith, S. N., Zhao, L., Wu, W., & Mobley, H. L. T. (2017). Genome-wide transposon mutagenesis of *Proteus mirabilis*: Essential genes, fitness factors for catheter-associated urinary tract infection, and the impact of polymicrobial infection on fitness requirements. *PLoS Pathog*, *13*(6), e1006434. https://doi.org/10.1371/journal.ppat.1006434

Blattner, F. R., Plunkett, G., 3rd, Bloch, C. A., Perna, N. T., Burland, V., Riley, M., Collado-Vides, J., Glasner, J. D., Rode, C. K., Mayhew, G. F., Gregor, J., Davis, N. W., Kirkpatrick, H. A., Goeden, M. A., Rose, D. J., Mau, B., & Shao, Y. (1997). The complete genome sequence of *Escherichia coli* K-12. *Science*, *277*(5331), 1453–1462. https://doi.org/10.1126/science.277.5331.1453

Boyd, E. F., Wang, F. S., Beltran, P., Plock, S. A., Nelson, K., & Selander, R. K. (1993). *Salmonella* reference collection B (SARB): strains of 37 serovars of subspecies I. *J Gen Microbiol*, *139 Pt 6*, 1125–1132. https://doi.org/10.1099/00221287-139-6-1125

Chamnongpol, S., & Groisman, E. A. (2002). $Mg^{2+}$ homeostasis and avoidance of metal toxicity. *Mol Microbiol*, *44*(2), 561–571. https://doi.org/10.1046/j.1365-2958.2002.02917.x

Chari, A., Oakeson, K. F., Enomoto, S., Jackson, D. G., Fisher, M. A., & Dale, C. (2015). Phenotypic characterization of *Sodalis praecaptivus* sp. nov., a close non-insect-associated member of the Sodalis-allied lineage of insect endosymbionts. *Int J Syst Evol Microbiol*, *65*(Pt 5), 1400–1405. https://doi.org/10.1099/ijs.0.000091

Daligault, H. E., Davenport, K. W., Minogue, T. D., Bishop-Lilly, K. A., Broomall, S. M., Bruce, D. C., Chain, P. S., Coyne, S. R., Frey, K. G., Gibbons, H. S., Jaissle, J., Koroleva, G. I., Ladner, J. T., Lo, C. C., Munk, C., Palacios, G. F., Redden, C. L., Rosenzweig, C. N., Scholz, M. B., & Johnson, S. L. (2014). Whole-genome assemblies of 56 burkholderia species. *Genome Announc*, *2*(6). https://doi.org/10.1128/genomeA.01106-14

Datsenko, K. A., & Wanner, B. L. (2000). One-step inactivation of chromosomal genes in *Escherichia coli* K-12 using PCR products. *Proc Natl Acad Sci U S A*, *97*(12), 6640–6645. https://doi.org/10.1073/pnas.120163297

Datta, S., Costantino, N., & Court, D. L. (2006). A set of recombineering plasmids for gram-negative bacteria. *Gene*, *379*, 109–115. https://doi.org/10.1016/j.gene.2006.04.018

Duchaud, E., Rusniok, C., Frangeul, L., Buchrieser, C., Givaudan, A., Taourit, S., Bocs, S., Boursaux-Eude, C., Chandler, M., Charles, J. F., Dassa, E., Derose, R., Derzelle, S., Freyssinet, G., Gaudriault, S., Medigue, C., Lanois, A., Powell, K., Siguier, P.,…Kunst, F. (2003). The genome sequence of the entomopathogenic bacterium *Photorhabdus luminescens*. *Nat Biotechnol*, *21*(11), 1307–1313. https://doi.org/10.1038/nbt886

Fields, P. I., Swanson, R. V., Haidaris, C. G., & Heffron, F. (1986). Mutants of *Salmonella typhimurium* that cannot survive within the macrophage are avirulent. *Proc Natl Acad Sci U S A*, *83*(14), 5189–5193. https://doi.org/10.1073/pnas.83.14.5189

Guzman, L. M., Belin, D., Carson, M. J., & Beckwith, J. (1995). Tight regulation, modulation, and high-level expression by vectors containing the arabinose PBAD promoter. *J Bacteriol*, *177*(14), 4121–4130. https://doi.org/10.1128/jb.177.14.4121-4130.1995

Jacobs, M. A., Alwood, A., Thaipisuttikul, I., Spencer, D., Haugen, E., Ernst, S., Will, O., Kaul, R., Raymond, C., Levy, R., Chun-Rong, L., Guenthner, D., Bovee, D., Olson, M. V., & Manoil, C. (2003). Comprehensive transposon mutant library of *Pseudomonas aeruginosa*. *Proc Natl Acad Sci U S A*, *100*(24), 14339–14344. https://doi.org/10.1073/pnas.2036282100

Keller, C.M., Kendra, C.G., Bruna, R.E., Craft, D., Pontes, M.H. (2021). Genetic Modification of *Sodalis* Species by DNA Transduction. *mSphere*. 6(1):e01331-20. https://doi.10.1128/mSphere.01331-20

Lee, E. J., Pontes, M. H., & Groisman, E. A. (2013). A bacterial virulence protein promotes pathogenicity by inhibiting the bacterium's own F1Fo ATP synthase. *Cell*, *154*(1), 146–156. https://doi.org/10.1016/j.cell.2013.06.004
Maloney, K. E., & Valvano, M. A. (2006). The *mgtC* gene of *Burkholderia cenocepacia* is required for growth under magnesium limitation conditions and intracellular survival in macrophages. *Infect Immun*, *74*(10), 5477–5486. https://doi.org/10.1128/IAI.00798-06
Pontes, M. H., & Groisman, E. A. (2018). Protein synthesis controls phosphate homeostasis. *Genes Dev*, *32*(1), 79–92. https://doi.org/10.1101/gad.309245.117
Pontes, M. H., & Groisman, E. A. (2019). Slow growth determines nonheritable antibiotic resistance in Salmonella enterica. *Sci Signal*, *12*(592). https://doi.org/10.1126/scisignal.aax3938
Pontes, M. H., Yeom, J., & Groisman, E. A. (2016). Reducing Ribosome Biosynthesis Promotes Translation during Low $Mg^{2+}$ Stress. *Mol Cell*, *64*(3), 480–492. https://doi.org/10.1016/j.molcel.2016.05.008
Ramage, B., Erolin, R., Held, K., Gasper, J., Weiss, E., Brittnacher, M., Gallagher, L., & Manoil, C. (2017). Comprehensive Arrayed Transposon Mutant Library of *Klebsiella pneumoniae* Outbreak Strain KPNIH1. *J Bacteriol*, *199*(20). https://doi.org/10.1128/JB.00352-17
Schlechter, R. O., Jun, H., Bernach, M., Oso, S., Boyd, E., Munoz-Lintz, D. A., Dobson, R. C. J., Remus, D. M., & Remus-Emsermann, M. N. P. (2018). Chromatic Bacteria - A Broad Host-Range Plasmid and Chromosomal Insertion Toolbox for Fluorescent Protein Expression in Bacteria. *Front Microbiol*, *9*, 3052. https://doi.org/10.3389/fmicb.2018.03052
Soncini, F. C., Vescovi, E. G., & Groisman, E. A. (1995). Transcriptional autoregulation of the *Salmonella typhimurium phoPQ* operon. *J Bacteriol*, *177*(15), 4364–4371. https://doi.org/10.1128/jb.177.15.4364-4371.1995
Staggs, T. M., & Perry, R. D. (1991). Identification and cloning of a fur regulatory gene in *Yersinia pestis*. *J Bacteriol*, *173*(2), 417–425. https://doi.org/10.1128/jb.173.2.417-425.1991
Stover, C. K., Pham, X. Q., Erwin, A. L., Mizoguchi, S. D., Warrener, P., Hickey, M. J., Brinkman, F. S., Hufnagle, W. O., Kowalik, D. J., Lagrou, M., Garber, R. L., Goltry, L., Tolentino, E., Westbrock-Wadman, S., Yuan, Y., Brody, L. L., Coulter, S. N., Folger, K. R., Kas, A.,…Olson, M. V. (2000). Complete genome sequence of *Pseudomonas aeruginosa* PAO1, an opportunistic pathogen. *Nature*, *406*(6799), 959–964. https://doi.org/10.1038/35023079
Thongdee, M., Gallagher, L. A., Schell, M., Dharakul, T., Songsivilai, S., & Manoil, C. (2008). Targeted mutagenesis of *Burkholderia thailandensis* and *Burkholderia pseudomallei* through natural transformation of PCR fragments. *Appl Environ Microbiol*, *74*(10), 2985–2989. https://doi.org/10.1128/AEM.00030-08
Tucker, A. T., Nowicki, E. M., Boll, J. M., Knauf, G. A., Burdis, N. C., Trent, M. S., & Davies, B. W. (2014). Defining gene-phenotype relationships in *Acinetobacter baumannii* through one-step chromosomal gene inactivation. *mBio*, *5*(4), e01313–01314. https://doi.org/10.1128/mBio.01313-14
Walker, K. A., Maltez, V. I., Hall, J. D., Vitko, N. P., & Miller, V. L. (2013). A phenotype at last: essential role for the *Yersinia enterocolitica* Ysa type III secretion system in a *Drosophila melanogaster* S2 cell model. *Infect Immun*, *81*(7), 2478–2487. https://doi.org/10.1128/IAI.01454-12
